# Evidence for microbiome-dependent chilling tolerance in sorghum

**DOI:** 10.1101/2021.03.02.433195

**Authors:** Sonya Erlandson, Raju Bheemanahalli, Nisarga Kodadinne Narayana, Marina Johnson, Christopher Graham, S. V. Krishna Jagadish, Senthil Subramanian

## Abstract

Early season chilling stress is a major constraint on sorghum production in temperate climates. Chilling-tolerant sorghum is an active area of development, but the potential for early season microbial-enhanced chilling tolerance in sorghum has not yet been explored. In this study, we characterized traits of field-grown sorghum accessions in response to chilling and non-chilling temperatures and the corresponding cohorts of phyllosphere fungal and bacterial taxa. Further, we characterized the effects of chilling temperatures and microbial inoculation on sorghum accession traits in a growth chamber experiment. By comparing sorghum trait responses under chilling stress with and without soil microbial inoculation, we were able to detect a potential microbe-dependent sorghum response to chilling stress. Four sorghum genotypes showed a negative response to chilling stress with vs. without microbial inoculation, while five sorghum accessions show increased shoot biomass or leaf area under chilling stress when inoculated with a soil microbiome. These differential responses provides opportunities to exploit beneficial microbial taxa for enhancing early-stage chilling tolerance in sorghum, with a potential to be extended to other crops.

## Introduction

Sorghum [(*Sorghum bicolor* L.) Moench] is one of the five major crop species in the world and is grown for grain or biomass. Diverse uses of sorghum for food, energy, paper, fertilizer, and feed makes it an economically important crop (Tari *et al*. 2013). However, as a plant that evolved in the African tropics, sorghum is sensitive to chilling temperatures, below 15°C (Yu and Tuinstra 2001). Temperature sensitivity has limited the geographic range of sorghum cultivation in colder regions, like the upper Midwest in the US, where it could otherwise be an attractive alternative crop for grain and biomass. In these colder regions, sorghum requires later planting due to early-stage (germination, emergence and seedling vigor) chilling sensitivity, which ultimately decreases biomass and yield (Maulana *et al*. 2013, Tarhi *et al*. 2013). A recent crop modelling exercise using sorghum planting dates from 1986-2015 in Kansas, recorded a 7 to 11% increase in yield with early planting, compared to normal and late planting (Raymundo *et al*. 2021). The study concluded that incorporating early-stage chilling tolerance in sorghum would allow for early planting, and hence a means to increase productivity extended growing season and by reducing the risk of terminal droughts.

While sorghum germplasm resources are diverse and under development for early-stage chilling tolerance (Chopra *et al*. 2015, Marla *et al*. 2019, Moghimi *et al*. 2019, Ostmeyer *et al*. 2020), the potential for early-stage microbial-enhanced chilling tolerance in sorghum has not yet been explored. Simultaneously evaluating diverse sorghum genotypes for genetic chilling tolerance and microbe-enhanced chilling tolerance would leverage a novel synergistic approach to early-stage chilling tolerance. Plant genotype-environment-microbe interactions in field conditions are highly complex (Bakker *et al*. 2020, Hohmann *et al*. 2020). Developing broadly effective microbial enhancements for plant abiotic stress will require scrutinizing plant genotype – microbe – environment interactions in controlled experiments.

Plant-associated microbes can improve abiotic stress tolerance in plants and exploiting these interactions promises new ways to improve crop production and agricultural practices (Backer *et al*. 2018, Lata *et al*. 2018). Various studies have addressed microbial impacts on plant chilling tolerance; focusing on potential mechanisms of a single microbe in conferring resilience against adverse conditions (Barka *et al*. 2006, Ding *et al*. 2011, Kang *et al*. 2015, Wang *et al*. 2015, Subramanian *et al*. 2016, Zhou *et al*. 2021). However, few studies have assessed changes in plant response to chilling associated with interactions at the microbial community level (Beirinckx *et al*. 2020, Cloutier *et al*. 2020). No studies, to our knowledge, have attempted to isolate the overall effects of the presence of a microbial community on the ability of seedlings to tolerate chilling stress. A basic understanding of how the presence of a diverse microbiome affects sorghum seedling vigor under chilling stress, and how the sorghum microbiome changes in response to chilling temperatures, is important context for further work on individual microbial isolates and mechanisms of chilling tolerance.

Prior studies have identified candidate genes and assessed agronomic outcomes for early-stage chilling tolerance in grain sorghum diversity panel (Moghimi *et al*. 2019). We utilized a subset of these sorghum accessions based on their field phenotypes (Bheemanahalli *et al*., 2019), to attempt to isolate the contribution of microbes to the plant chilling stress response and compare the microbial contribution across accessions. We hypothesized that grain sorghum accessions will exhibit diverse phenotypic responses to chilling stress, and that these diverse responses to chilling stress can be broken down into accession-specific component (microbe-independent chilling tolerance) and an accession-microbe interaction component (microbe-dependent chilling tolerance). We characterized field-grown sorghum accessions traits, leaf fungal, and leaf bacterial communities in response to chilling and non-chilling temperatures. Microbial community composition was significantly impacted by geographical location and temperature while the sorghum accessions had little to no effect. We further tested the effects of chilling temperatures and microbial inoculation on sorghum performance in a growth chamber experiment. By comparing sorghum traits under chilling stress with and without microbial inoculation in a growth chamber experiment, we detected a microbe-dependent sorghum response to chilling stress. Further, these results were compared to multi-environment field studies to provide a realistic context for interpreting the microbe-dependent chilling response.

## Methods

### Field experiment design

Twenty sorghum accessions (15 tolerant and 5 sensitive selected from the sorghum association panel of 400 accessions based on field trials in 2018) were grown at Kansas State University, Manhattan, Kansas and South Dakota State University West River Research Farm, Sturgis, South Dakota. Planting was conducted in two intervals as early and regular planting in 2019. Based on the sorghum production practices at each site, regular planting dates were June 11 in Kansas and June 5 in South Dakota. The early planting date was April 24 in Kansas and May 16 in South Dakota. The early planting date was intended to induce chilling stress temperatures and regular planting date as the optimal sorghum growing temperatures. Early planting dates were selected based on historic average air temperatures at the respective field sites. The target air temperature for early planting (chilling stress) was 13°C and that for regular planting (control) was 20°C. The experiment was laid out in a split-plot randomized complete block design with three replications. Each genotype was planted in four (Kansas) or two (South Dakota) row plots, and replicated thrice for each planting type (temperature treatment). Each row was 3.6 m (12 ft) long and accommodated 48 seeds (Kansas) or 7.3m (24 ft) long and accommodated 101 seeds (South Dakota) with an inter-row spacing of 0.75 m. At the Kansas site, the maximum/minimum soil temperature at ~10 cm depth for different planting dates was recorded at 15 min intervals using soil temperature data loggers (Onset Computer Corporation, Bourne, MA, USA). Air and bare soil (4-inch depth) temperatures for the South Dakota West River Research Farm were obtained from the Mesonet weather station at the site (South Dakota Mesonet, https://mesonet.sdstate.edu/archive). Minimum and maximum air temperatures for each day were averaged (Supplementary Fig. 1 and 2).

**Figure 1.**
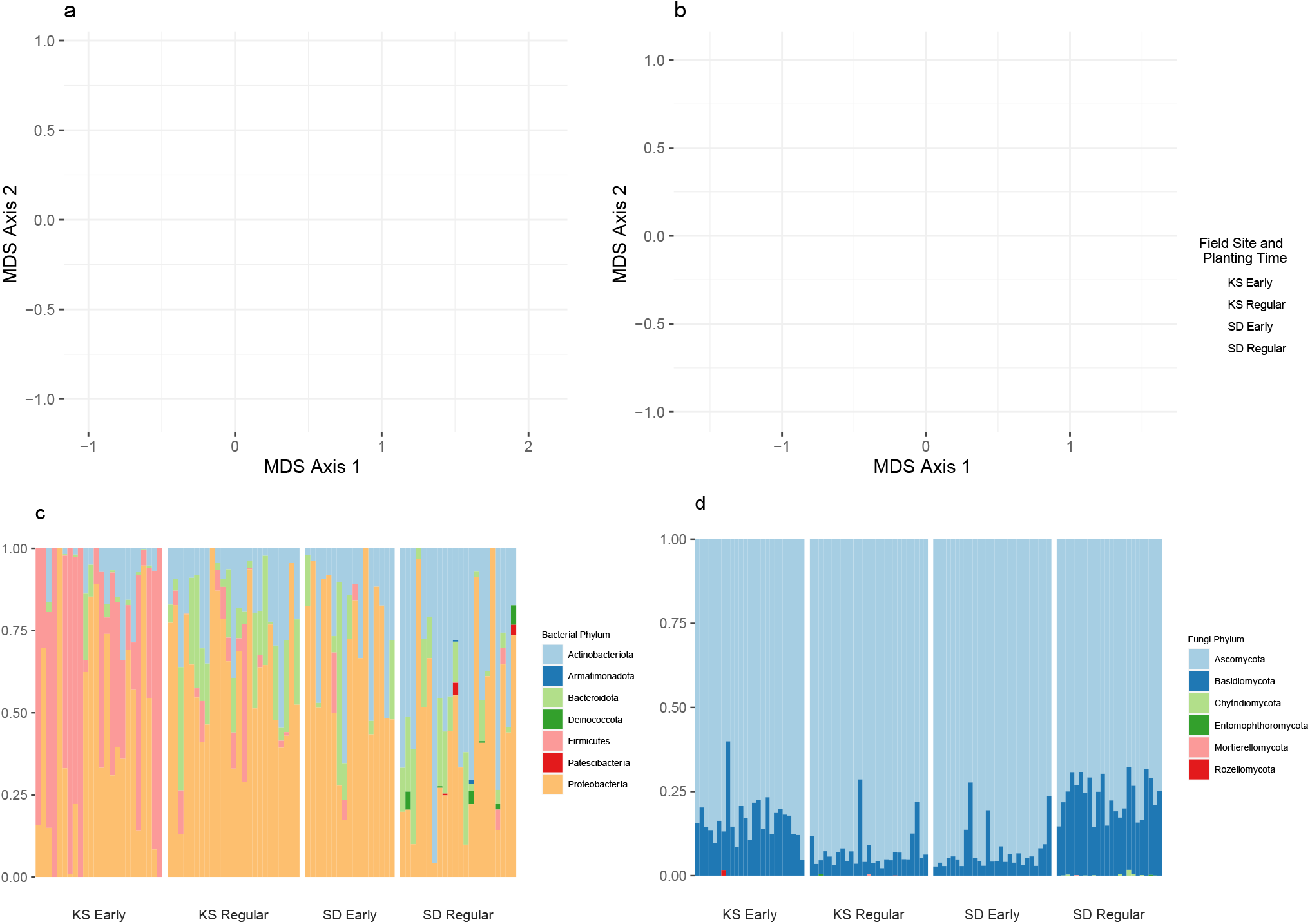
Principle coordinates analysis (multidimensional scaling) of bacterial (a) and fungal (b) communities. Points represent ASV communities of sorghum leaf samples and are colored by site and time of collection Kansas (KS), South Dakota (SD), early planting (Early) and regular planting (Regular). Bacterial ASVs dissimilarity in samples from regular Kansas (KS) planting time separate clearly from other samples along MDS1, while MDS2 shows community differences between Kansas early planting and South Dakota samples. Fungal community dissimilarity clearly groups samples by site and planting time. Barplots (c) and (d) show relative abundance of bacterial (c) and fungal (d) phyla across field site and time.

**Figure 2.**
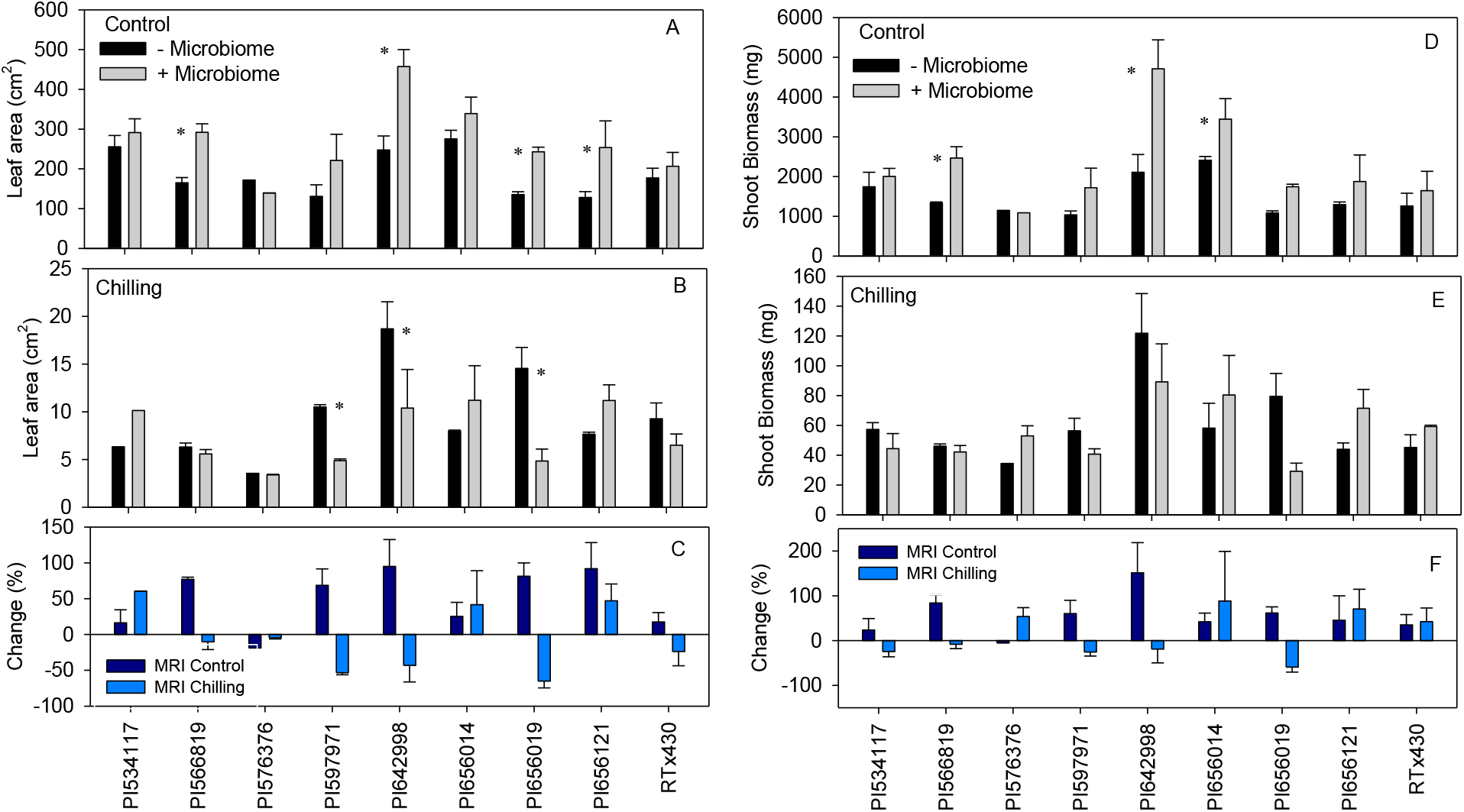
Leaf area (A-C) and shoot biomass (D-F) of sorghum accessions at 45 DAP with and without microbe inoculation under control (A and D; 30°C/20°C, day/night) and chilling stress (B and E; 20°C/10°C, day/night). C and F Microbiome Response Index (MRI) indicates the percentage change with and without microbe inoculation under control and chilling conditions. * represents a significant difference between treatments (microbiome and accession) at 5%.

### Field measurements

Emergence percentage, leaf area, and shoot biomass per seedling were measured under both early planting and regular planting conditions. Seedling emergence percentage was calculated as the ratio of the total number of emerged seedlings to the total number of planted seeds, multiplied by 100. Destructive sampling was done 45 days after planting (DAP) for both planting conditions. Leaf area was estimated by a leaf area meter (LI-3100 area meter, LI-COR. Inc., Lincoln, NE, USA) before oven drying. The shoot was separately and oven-dried at 60°C for 72 h, and shoot biomass (g plant^−1^) was recorded after reaching constant weight to obtain shoot biomass per seedling. Shoot biomass per row was calculated as a product of average seedling shoot biomass x emergence count per row.

### Field experiment leaf microbiome characterization

A subset of accessions were selected for microbial analysis (Table 1, same accessions used for growth chamber experiment). The second fully expanded leaf was collected from two representative plants per replicate. Leaves were stored at 4°C for up to 24 hours and frozen at −80°C until DNA extraction. DNA in leaf tissue was purified with the Qiagen PowerSoil DNA Extraction Kit following manufacturer protocol, using equal mass from the two representative leaves per plant (0.12g + 0.12g = 0.24 g leaf tissue). Leaf tissue was crushed to powder with liquid N_2_ in a mortar and pestle before loading PowerSoil tube filled with C1 solution. Bacteria and fungi were characterized with frame-shifting, locus-specific primer sets with Illumina overhang adapters as in Lundberg *et al*. 2013. Briefly, four versions of each primer were generated, each containing a different number of nucleotides between the adapter sequence and locus-specific primer, then pooled equimolarly for locus-specific PCR. Bacterial primers 799F (Chelius and Triplett 2001) and 1115R (Wallace *et al*. 2018a, Wallace *et al*. 2018b) were used to amplify 16S V5-V6. Fungal primers ITS1f (Gardes and Bruns, 1993) and ITS2 (White *et al*. 1990) were used to amplify ITS1. Locus-specific PCR was followed by index PCR with Nextera Kit v2 sets A, B, C, D (see supplementary methods for primer design and thermocycler conditions). All PCR products were cleaned with HighPrep PCR Clean-up System (MagBio) and cleaned products were quantified with Qubit Broad Range DNA Assay (Thermo Fisher Scientific). Bacterial and fungal amplicons were pooled equimolarly and sequenced on an Illumina MiSeq 2 ⨯ 300 at South Dakota State University Genomics Sequencing Center following manufacturer protocol.

**Table 1.** Kansas. Emergence percentage, leaf area, and shoot biomass of sorghum accessions grown in the Kansas field site. Significance ranged between < 0.05 and < 0.001 probability level; NS, non-significant. Accessions with different letters are significantly different at P < 0.05 within respective planting conditions. “−”: no seedling emergence. Accessions with underlines are those selected for microbiome experiment.

| Accession | Emergence (%) |  |  | Leaf area (cm<sup>2</sup>) |  |  | Shoot biomass (g) |  |  | Shoot biomass per row (g) |  |  |
| --- | --- | --- | --- | --- | --- | --- | --- | --- | --- | --- | --- | --- |
|  | Regular | Early | % change | Regular | Early | % change | Regular | Early | % change | Regular | Early | % change |
| PI533755 | 14.58cde | 20.62efg | -41.43 | 1296.29d | 156.86ab | 87.90 | 29.19abcd | 0.81a | 97.23 | 202.31cde | 9.74bc | 95.19 |
| PI533831 | 4.17e | 6.87g | -64.75 | 2532.90bc | 58.40b | 97.69 | 20.27d | 0.29a | 98.57 | 40.73e | 1.05c | 97.42 |
| PI540816 | 9.11de | - |  | 2672.02bc | - |  | 40.86a | - |  | 194.37cde | - |  |
| PI655978 | - | - |  | - | - |  | - | - |  | - | - |  |
| PI655996 | - | - |  | - | - |  | - | - |  | - | - |  |
| <u>PI534117</u> | 16.49bcd | 54.25abc | -228.99 | 1914.25cd | 156.12ab | 91.84 | 36.81ab | 0.85a | 97.69 | 294.89bcd | 22.46ab | 92.38 |
| <u>PI566819</u> | 16.32bcd | 31.11cdef | -90.63 | 3097.40ab | 118.14ab | 96.19 | 34.49abc | 0.45a | 98.70 | 310.12bcd | 8.18bc | 97.36 |
| <u>PI576376</u> | 18.58bcd | 62.57a | -236.76 | 2474.75bc | 167.77a | 93.22 | 26.27abcd | 0.83a | 96.84 | 238.12bcde | 26.16a | 89.01 |
| PI576394 | 11.46de | 28.57defg | -149.30 | 2413.82bc | 139.00ab | 94.24 | 29.08abcd | 0.77a | 97.35 | 156.82de | 11.36abc | 92.76 |
| <u>PI597971</u> | 23.96abc | 19.17fg | 19.99 | 2591.99bc | 95.40ab | 96.32 | 34.02abcd | 0.53a | 98.44 | 383.44bc | 4.95c | 98.71 |
| PI642992 | 18.06bcd | 38.34bcdef | -112.29 | 3099.80ab | 93.43ab | 96.99 | 26.93abcd | 0.48a | 98.22 | 232.93bcde | 9.23bc | 96.04 |
| <u>PI642998</u> | 8.85de | 54.62abc | -517.18 | 2572.24bc | 151.84ab | 94.10 | 36.68ab | 0.84a | 97.71 | 179.52cde | 25.51a | 85.79 |
| PI651496 | 19.10bcd | 44.49abcde | -132.93 | 3613.29a | 97.58ab | 97.30 | 36.06ab | 0.54a | 98.50 | 319.18bcd | 11.69abc | 96.34 |
| PI655991 | 17.19bcd | 36.89bcdef | -114.60 | 3065.34ab | 103.78ab | 96.61 | 36.30ab | 0.51a | 98.60 | 300.15bcd | 9.32bc | 96.89 |
| <u>PI656014</u> | 22.92bc | 42.32abcdef | -84.64 | 2024.61cd | 125.46ab | 93.80 | 22.14cd | 0.63a | 97.15 | 239.06bcde | 14.22abc | 94.05 |
| <u>PI656019</u> | 23.96abc | 56.79ab | -137.02 | 2593.99bc | 114.10ab | 95.60 | 36.10ab | 0.63a | 98.25 | 425.29ab | 19.29abc | 95.46 |
| PI656023 | 19.44bcd | 42.68abcdef | -119.55 | 3103.51ab | 80.44b | 97.41 | 38.80ab | 0.48a | 98.76 | 361.51bcd | 9.87bc | 97.27 |
| PI656035 | 34.03a | 21.34efg | 37.29 | 2973.10ab | 91.58ab | 96.92 | 37.26ab | 0.45a | 98.79 | 600.80a | 4.66c | 99.22 |
| PI656038 | 16.15bcd | 49.91abcd | -209.04 | 2418.47bc | 121.33ab | 94.98 | 25.44bcd | 0.62a | 97.56 | 192.67ede | 15.49abc | 91.96 |
| <u>PI656121</u> | 26.39ab | 22.79efg | 13.64 | 2064.30cd | 90.81ab | 95.60 | 28.59abcd | 0.51a | 98.22 | 379.74bc | 5.57c | 98.53 |
| Treatment (T) | NS |  |  | < 0.001 |  |  | < 0.001 |  |  | < 0.01 |  |  |
| Accession (A) | < 0.001 |  |  | < 0.01 |  |  | NS |  |  | < 0.01 |  |  |
| T x A | < 0.001 |  |  | < 0.001 |  |  | NS |  |  | < 0.01 |  |  |

### Amplicon bioinformatics

Demultiplexed Illumina reads were evaluated for sequencing quality with FastQC (v0.11.8). Each primer was trimmed from reads using Cutadapt (v2.0) and quality checked again with FastQC to verify removal of PCR primers. Trimmed reads were filtered, quality trimmed, denoised (error corrected), merged, chimeric sequences removed, and taxonomy assigned using dada2 (v1.10.1) in R (v3.6.1). Dada2 parameters for filtering ITS reads were maximum N of 0, removing bases with quality below 2, a minimum read length of 50 bp, and a maximum expected error of 2 for both read pairs. and fungal taxonomy assigned using the UNITE database (vFeb2020) and RDP classifier implemented in dada2. Parameters for filtering 16S reads in dada2 were the same as fungi, with the exception of truncating read length to 200. Bacteria taxonomy was assigned with SILVA SSU database (v138) and the RDP classifier in dada2. Amplicon sequence variant (ASV) (similar to 100% Operational Taxonomic Unit) tables were filtered prior to statistical analysis, 16S and ITS reads were analyzed independently. Taxonomic assignment was used to remove reads only classified to kingdom for both 16S and ITS reads (i.e. taxonomy had to be assigned to at least phylum level). On manual inspection, it was found that reads classified only to kingdom tended to be sorghum DNA. Reads that matched to mitochondria or chloroplasts were removed from the 16S data set. Reads present in 16S negative controls were also removed. ASVs with a read abundance less than 0.001% of the whole data set were removed and samples with fewer than 15 (16S) or 1000 (ITS) reads were removed.

Rarefaction curves were calculated for ASV tables before and after filtering to determine if sampling (sequencing) depth saturated the number of ASVs in the plant sample.

### Growth chamber study

#### Plant material and growth conditions

In this study, chilling (20/10°C, day/night) and control (30/20°C, day/night) temperature treatments were used to create temperature conditions similar to early and regular planting conditions in the field experiment. Three seeds of similar size for each of the eight-selected sorghum accessions (five high biomass and three low biomass accessions [Table 1], one sensitive genotype, RTx430) were sown at a depth of 2-3 cm in a pot (10 cm×10 cm×30 cm, square pot) filled with 150 g of 3:1 proportion of double-autoclaved vermiculite: perlite mix. The pots were placed in four growth chambers, with three pots maintained per accession in each chamber, totaling to 24 replicate pots across treatments (2 inoculum x 2 temperature treatments). The chambers were programmed to reach the daytime (08:00 h to 17:00 h) target temperature, following a gradual increase from 20°C to 30°C (control) and from 10°C to 20°C (chilling stress) over 3 h (05:00 h to 08:00 h). A similar gradual transition was followed from the maximum daytime temperature [30°C (control) or 20°C (chilling)] to night-time temperature (20°C or 10°C, respectively) over a 3 h (17:00 h and 20:00 h) period. Both control and chilling stress chambers were maintained at a 12 h photoperiod, with 800 μmol m^−2^ s^−1^ light intensity at 5 cm above the canopy and 60% relative humidity (RH). Air temperature and RH were recorded every 15 min using HOBO UX 100-003 temperature/RH data loggers (Onset Computer Corp., Bourne, MA, USA) in all growth chambers. The pots were watered throughout the experiment as required using distilled water.

#### Soil microbe extraction and treatments

Two potting mixes that were identical except for their microbial communities were used in the study. Microbial cells were extracted following the soil wash method as described by Wagner *et al*. (2014). In brief, six soil subsamples representing ~1 acre were collected at a depth of ~10-30 cm from the same Manhattan, KS field used for the field experiment. Collected samples were homogenized by hand after removing rocks and other large debris. The homogenized soil samples were sieved through 3.5 mm wire mesh before microbial extraction. A soil subsample of 75 g was stirred into 1 L of 2.5 mM MES monohydrate (Sigma Aldrich, St. Louis, MO, USA) in sterile distilled water (diH_2_O) (pH adjusted to 5.7 with KOH). The soil suspensions were vacuum filtered to remove particulates after 30 min. Filtrates were centrifuged for 20 min, 4000 rpm at 25°C to pellet down microbial cells. Distilled water and MES solutions were sterilized via autoclave. Supernatants were discarded and the microbe-enriched pellets were re-suspended in sterile 2.5 mM MES solution. Pots were saturated with sterile distilled water (1 L; free from microbial contamination) one day before planting sorghum seeds. One day post-planting, each microbial inoculation treatment pot received 50 mL diluted inoculum solution (100 mL of microbial suspension and 900 mL of sterile diH_2_O). An additional 5 mL of undiluted microbial suspension was pipetted into each microbial inoculation pot (top-dressed). For the non-inoculated control, pots were saturated with an inoculum-free solution containing 100 mL sterile 2.5 mM MES and 900 mL of sterile diH_2_O. An additional 5 mL sterile 2.5 mM MES was pipetted into each pot (top-dressed). Treatments derived by this process are termed “+ Microbiome” and “−Microbiome” to indicate microbial inoculation and no microbial inoculation, respectively.

#### Growth chamber experiment design

Chlorophyll index (Model 502, Spectrum Technologies, IL, USA) and effective quantum yield (QY) of photosystem II (FluorPen FP 100, Photon System Instruments, Ltd., Brno, Czech Republic) were measured on the topmost fully expanded leaf before harvest. The seedlings of all the accessions were harvested exactly 45 DAP. The total leaf area was measured using a leaf area meter (LI-3100 area meter, LI-COR, Lincoln, NE, USA). The shoot dry weight was recorded after oven drying at 60°C.

#### Estimation of indices to explore microbiome dependent and independent effect

Different indices were calculated to determine the microbiome influence on various traits measured under control and chilling using the following five different equations. The Microbiome Response Index (MRI, Control or Chilling) was calculated using equations 1 and 2. Trait values in the microbial inoculation treatment are expressed as a percentage of that in control treatment. A positive value indicates a benefit from the microbial inoculation and *vice versa*. Equations 1 and 2 identify accessions that can benefit from microbial inoculation under control and chilling, respectively. Chilling Tolerance Index without microbe inoculation (CTIm; Equation 3) was used to evaluate the microbiome-independent effect under chilling tolerance of each accession for a given trait. Chilling Tolerance Index with microbe inoculation (CTIM; Equation 4) represents the combined effect of microbiome-dependent and microbiome-independent chilling tolerance of each accession. Therefore, equation 5 (CTIM minus CTIm) values indicate the level of microbiome-dependent chilling tolerance for each trait. Accessions with positive values for a given trait are those that indicate an enhanced microbiome-dependent chilling tolerance.

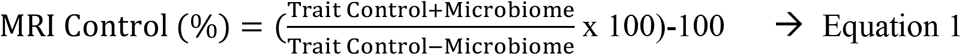

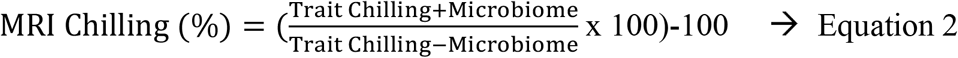

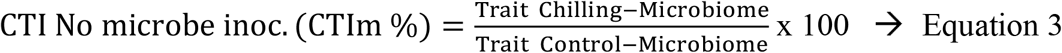

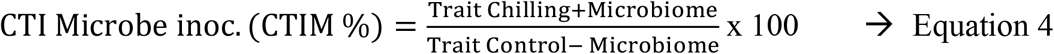

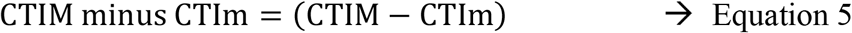

### Statistical analysis

**Field**The experimental layout was a split-plot with the randomized complete block design, with three replications. An ANOVA was performed for seedling emergence percentage, leaf area, and shoot biomass to estimate the significant difference between early and regular planting using the library (agricolae) in RStudio 3.6.1 (https://rstudio.com/). Means were separated using the least significant difference (LSD) when treatments and interactions were significant at P≤0.05. The same set of analyses were conducted in JMP (SAS Institute, Inc.) or R (with the same methods as Kansas) for South Dakota field data.

Analyses for 16S and ITS amplicon data were conducted separately. ASV community data tables were transformed by weighted centered log-ratio with the easyCODA package for community analyses. Main factors and their interactions (site, planting time, accession) were assessed for significance in explaining differences in bacterial and fungal leaf-associated community composition with permutational ANOVAs (adonis in vegan). Microbial community beta-diversity was visualized with principal coordinates analysis (capscale in vegan). Wald tests on coefficients from negative binomial generalized linear models were used to identify differentially abundant taxa between early and regular planting with the DESeq2 package. Taxa were investigated at the phylum and genus levels and were considered significantly different at an adjusted P≤0.05 and log2 Fold Change < or > 0. Sample-wise Shannon index was calculated on rarefied datasets. Differences in mean Shannon index between sites and planting times were tested with ANOVA and when significant at P≤0.05 separated with Tukey’s HSD. All microbial statistical analyses were completed in RStudio v1.2.5019 with R v 3.6.1.

### Growth chamber

An ANOVA was performed for all traits to estimate the impact of the microbiome (+/-) of control and chilling using the library (agricolae) in RStudio 3.6.1 (https://rstudio.com/). A split-plot with the completely randomized design was followed to analyze the data, considering the microbiome as the main plot, accessions as the subplot under each temperature treatment. Means were separated using the least significant difference (LSD) when treatments and interactions were significant at P≤0.05.

## Results

### Field evaluation of sorghum accessions identified those with consistent chilling tolerance phenotypes in both KS and SD locations

The goal here was to evaluate a set of twenty sorghum accessions with varying levels of chilling tolerance at KS and SD locations to identify those that show consistent chilling response phenotypes at both locations. Early planting soil and air temperatures were lower by 8°C and 7°C, respectively, compared to regular planting conditions in Kansas (Fig. S1). In South Dakota, early planting soil and air temperatures were lower by 9°C and 8°C compared to regular planting (Fig. S2). Early planting had a significant (P<0.01) negative impact on leaf area, shoot biomass per seedling, and shoot biomass per row (Table 1). In contrast, the lack of a significant effect of chilling stress on the emergence percentage was observed (Table 1). In South Dakota, early planting had a significant (P <0.01) negative impact on emergence percentage, leaf area, shoot biomass per seedling, and shoot biomass per row (Table 2). Emergence in the early planting treatment ranged from 6% to 62% and 4% to 34% in the regular planting conditions in Kansas. A similar trend was found in South Dakota, with early planting emergence ranged between 0% to 66% and 0% to 60% emergence in regular planting (Table 2). Under early planting conditions, the emergence percentage of PI576376 was significantly greater than PI533831, where all other accessions were statistically the same at the Kansas field site (Table 1). In South Dakota, accession PI656014 had emergence percentage significantly higher than all other accessions (Table 2). Accessions showed significant difference for leaf area across sites. Leaf area due to chilling stress conditions was reduced by >90%, averaged across all the accessions in Kansas and between 70% to 91% in South Dakota (Table 1, 2). Similar to emergence percentage, the leaf area of PI576376 was significantly greater than PI533831 under early planting conditions in Kansas. The leaf area of PI656014, PI576376, and PI642998 were significantly higher than other accessions in South Dakota (Table 2). Seedling shoot biomass was significantly (P<0.001) affected by chilling stress at both sites. However, there was no significant difference in shoot biomass per seedling among sorghum accessionsin either site. On average, seedling shoot biomass across accessions was reduced by 96% to 98% with early planting over regular planting in Kansas and 78% to 92% in South Dakota (Table 1, 2). However, shoot biomass per row of PI576376 and PI642998 was significantly greater than nine of the other accessions under early planting conditions, in Kansas (PI566819, PI533755, PI533831, PI597971, PI642992, PI655991, PI656023, PI656035, PI656121, Table 1). This suggests that the combined effect of reduction in emergence percentage and shoot biomass per seedling led to overall reduction in biomass yields in some accessions. In early planting in South Dakota, PI656014 had significantly greater shoot biomass per row than PI655991 and PI597971 (Table 2). All field physiological measurements showed a significant treatment x accession interaction except shoot biomass per seedling in Kansas (Table 1, Table 2).

**Table 2.**
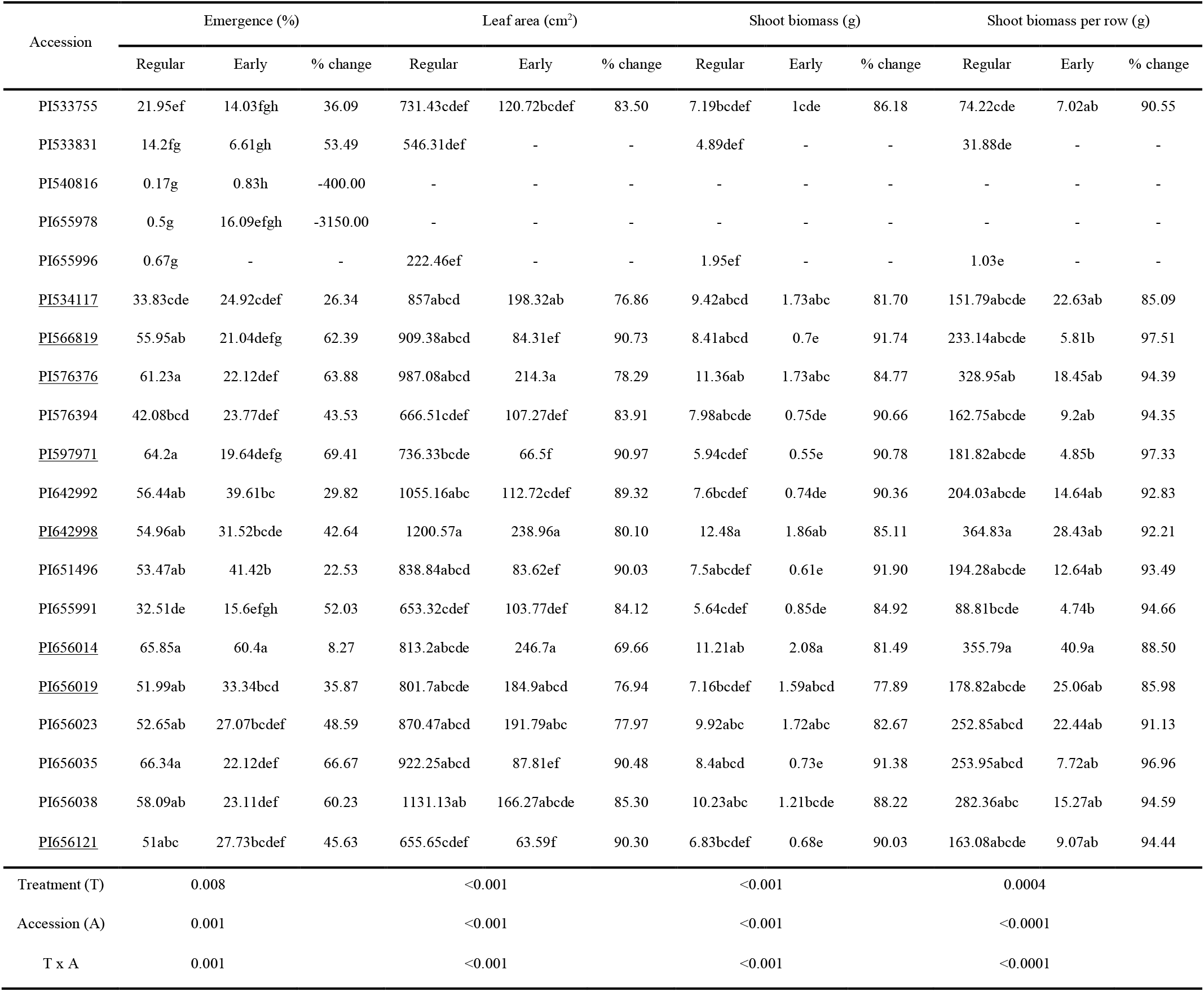
South Dakota. Emergence percentage, leaf area, and shoot biomass of sorghum accessions grown in the South Dakota field site. Accessions with different letters are significantly different at P = 0.05 within respective planting conditions. “-”: no seedling emergence. Accessions with underlines are those selected for microbiome experiment.

| Accession | Emergence (%) |  |  | Leaf area (cm<sup>2</sup>) |  |  | Shoot biomass (g) |  |  | Shoot biomass per row (g) |  |  |
| --- | --- | --- | --- | --- | --- | --- | --- | --- | --- | --- | --- | --- |
|  | Regular | Early | % change | Regular | Early | % change | Regular | Early | % change | Regular | Early | % change |
| PI533755 | 21.95ef | 14.03fgh | 36.09 | 731.43cdef | 120.72bcdef | 83.50 | 7.19bcdef | 1cde | 86.18 | 74.22cde | 7.02ab | 90.55 |
| PI533831 | 14.2fg | 6.61gh | 53.49 | 546.31def | - | - | 4.89def | - | - | 31.88de | - | - |
| PI540816 | 0.17g | 0.83h | -400.00 | - | - | - | - | - | - | - | - | - |
| PI655978 | 0.5g | 16.09efgh | -3150.00 | - | - | - | - | - | - | - | - | - |
| PI655996 | 0.67g | - | - | 222.46ef | - | - | 1.95ef | - | - | 1.03e | - | - |
| <u>PI534117</u> | 33.83cde | 24.92cdef | 26.34 | 857abcd | 198.32ab | 76.86 | 9.42abcd | 1.73abc | 81.70 | 151.79abcde | 22.63ab | 85.09 |
| <u>PI566819</u> | 55.95ab | 21.04defg | 62.39 | 909.38abcd | 84.31ef | 90.73 | 8.41abcd | 0.7e | 91.74 | 233.14abcde | 5.81b | 97.51 |
| <u>PI576376</u> | 61.23a | 22.12def | 63.88 | 987.08abcd | 214.3a | 78.29 | 11.36ab | 1.73abc | 84.77 | 328.95ab | 18.45ab | 94.39 |
| PI576394 | 42.08bcd | 23.77def | 43.53 | 666.51cdef | 107.27def | 83.91 | 7.98abcde | 0.75de | 90.66 | 162.75abcde | 9.2ab | 94.35 |
| <u>PI597971</u> | 64.2a | 19.64defg | 69.41 | 736.33bcde | 66.5f | 90.97 | 5.94cdef | 0.55e | 90.78 | 181.82abcde | 4.85b | 97.33 |
| PI642992 | 56.44ab | 39.61bc | 29.82 | 1055.16abc | 112.72cdef | 89.32 | 7.6bcdef | 0.74de | 90.36 | 204.03abcde | 14.64ab | 92.83 |
| <u>PI642998</u> | 54.96ab | 31.52bcde | 42.64 | 1200.57a | 238.96a | 80.10 | 12.48a | 1.86ab | 85.11 | 364.83a | 28.43ab | 92.21 |
| PI651496 | 53.47ab | 41.42b | 22.53 | 838.84abcd | 83.62ef | 90.03 | 7.5abcdef | 0.61e | 91.90 | 194.28abcde | 12.64ab | 93.49 |
| PI655991 | 32.51de | 15.6efgh | 52.03 | 653.32cdef | 103.77def | 84.12 | 5.64cdef | 0.85de | 84.92 | 88.81bcde | 4.74b | 94.66 |
| <u>PI656014</u> | 65.85a | 60.4a | 8.27 | 813.2abcde | 246.7a | 69.66 | 11.21ab | 2.08a | 81.49 | 355.79a | 40.9a | 88.50 |
| <u>PI656019</u> | 51.99ab | 33.34bcd | 35.87 | 801.7abcde | 184.9abcd | 76.94 | 7.16bcdef | 1.59abcd | 77.89 | 178.82abcde | 25.06ab | 85.98 |
| PI656023 | 52.65ab | 27.07bcdef | 48.59 | 870.47abcd | 191.79abc | 77.97 | 9.92abc | 1.72abc | 82.67 | 252.85abcd | 22.44ab | 91.13 |
| PI656035 | 66.34a | 22.12def | 66.67 | 922.25abcd | 87.81ef | 90.48 | 8.4abcd | 0.73e | 91.38 | 253.95abcd | 7.72ab | 96.96 |
| PI656038 | 58.09ab | 23.11def | 60.23 | 1131.13ab | 166.27abcde | 85.30 | 10.23abc | 1.21bcde | 88.22 | 282.36abc | 15.27ab | 94.59 |
| <u>PI656121</u> | 51abc | 27.73bcdef | 45.63 | 655.65cdef | 63.59f | 90.30 | 6.83bcdef | 0.68e | 90.03 | 163.08abcde | 9.07ab | 94.44 |
| Treatment (T) | 0.008 |  |  | <0.001 |  |  | <0.001 |  |  | 0.0004 |  |  |
| Accession (A) | 0.001 |  |  | <0.001 |  |  | <0.001 |  |  | <0.0001 |  |  |
| T x A | 0.001 |  |  | <0.001 |  |  | <0.001 |  |  | <0.0001 |  |  |

When accessions were ranked based on shoot biomass per row, five accessions (PI534117, PI642998, PI656014, PI565019 and PI576376) consistently ranked among the top six at both Kansas and South Dakota. Similarly, PI656121, PI597971, and PI566819 performed consistently poorly at both Kansas and South Dakota, recording lower shoot biomass per row. These eight accessions were evaluated for leaf associated microbial communities to determine if and what impact planting site (KS vs SD), plating time (early – chilling stress vs. regular – optimal), and genotype (chilling tolerance) had on microbial community diversity and composition (Table 1 and 2).

### Site and planting time explain differences in leaf-associated microbial communities

Microbial communities from second fully opened leaves were evaluated by high throughput sequencing of bacterial 16S and fungal ITS amplicons. The sequencing run yielded 71,140,584 read pairs. After all filtering steps, 79,098 bacterial and 4,023,465 fungal reads were retained. Reads comprised 148 bacterial amplicon sequence variants (ASVs) in 88 samples and 1450 fungal ASVs in 103 samples. Rarefaction curves indicated that the sorghum leaf communities were well-sampled (Fig. S3).

We aimed to characterize and compare the bacterial and fungal taxa of the grain sorghum phyllosphere grown in early (chilling stress) and in regular (optimal) planting at the two locations. Common ASVs across all samples and ASVs unique to each site varied between fungi and bacteria. South Dakota and Kansas field sites share 41.9% bacterial ASVs (62) and 29.1% fungal ASVs (422). Bacterial ASVs unique to site were 28.4% for South Dakota (42) and 29.7% for Kansas (44). Fungal ASVs unique to site were 33% for South Dakota (479) and 37.9% for Kansas (549). The bacterial ASVs *Pseudomonas, Microbacterium*, and *Acidovorax* spp. were the three most abundant overall, present in 30, 25 and 19 samples out of 88 total at relative abundances of 13.5%, 10%, and 9.9%, respectively (Table S1). Fungal ASVs with highest relative abundance across the dataset were *Alternaria* sp. (21.3%), *Mycosphaerella tassiana* (15.9%), *Epicoccum dendrobii* (10.3%), which were present in all samples (Table S2). We also characterized relative abundance at each planting time within site. *Exiguobacterium* (30%), *Massilia* (24%), and *Paenibacillus* (13%) were genera with the highest relative abundance at early planting in Kansas, while *Pantoea* (42%), *Buchnera* (10%), and *Hymenobactr* (10%) had the highest relative abundance at early planting in South Dakota (Table S1). For regular planting samples, *Pseudomonas* (26%), *Microbacterium* (23%), *Acidovorax* (20%) were the three genera with the highest relative abundance in Kansas and *Microbacterium* (20%), *Rathayibacter* (16%), *Spirosoma* (10%) dominated in South Dakota (Table S1). The most abundant fungal taxa were more similar across all samples than bacteria. Fungal genera with highest relative abundance at early planting in Kansas were *Alternaria* (22%), *Mycosphaerella* (13%), and *Leptospora* (13%), and *Mycosphaerella* (39%), *Alternaria* (22%), *Epicoccum* (12%) in South Dakota (Table S2). In regular planting samples, *Alternaria* (44%), *Epicoccum* (14%), *Cladosporium* (11%) genera had highest relative abundance in Kansas and *Alternaria* (17%), *Epicoccum* (16%), *Mycosphaerella* (15%) in South Dakota (Table S2).

Next we determined how field site, planting time, and sorghum accession explained differences in the leaf-associated microbial community compositions. Multivariate ANOVAs indicated that field site, planting time, and field site x planting time interaction significantly explained variation in both microbial communities (Table S3, Fig. 1a,b). Sorghum accession accounted for 6.6% and 2.4% of variation in bacteria and fungi communities, respectively, but was not a significant predictor (Table S3). Site explained the most variation for fungal community composition (fungi = 48.8%, bacteria = 14.6%), while planting time explained the most variation for bacteria (bacteria = 19.7%, fungi = 16.4%) (Table S3). The microbial community differences between early and regular planting times depended on site, indicated by the significant site x planting time interaction and visualized in principle coordinates analysis plots (Table S3, Fig. 1). Bacterial communities form clear groups for Kansas early and regular planting times, but South Dakota planting time communities are less differentiated (Fig. 1). Fungal communities form four clear groups corresponding to the two sites and two planting times (Fig. 1). We also investigated how diversity changes across site and planting time. At the Kansas site, bacterial diversity was the same in early planting (mean H’ = 1.47 ±0.13) and regular planting samples (mean H’ = 1.54 ±0.17), while fungal diversity was higher in early planting than regular planting samples (mean H’ = 3.58 ±0.05 vs. 2.61 ±0.06, P<0.001). For South Dakota samples, both bacterial and fungal diversity were lower in early planting than regular planting samples (bacteria: mean H’ = 1.37 ±0.19 vs. 2.01 ±0.18, P=0.01, fungi: mean H’ = 2.61 ±0.07 vs. 3.34 ±0.07, P<0.001, Figs. S4, S5).

Finally, we identified individual genera and phyla that significantly changed in abundance between the two planting times. Differential abundance analysis between early planting and regular planting by site identified 19 bacterial genera in South Dakota and 24 in Kansas samples, with 4 genera more abundant in early planting South Dakota samples and 13 in Kansas (Fig. S6). Bacteria genera that showed consistent differential abundance at both sites (both relatively more abundant in early planting, for example) included *Brevundimonas* and *Pantoea* with higher abundance in early relative to regular planting and *Microbacterium, Pseudomonas*, and *Mucilaginibacter* with higher abundance in regular relative to early planting. Several bacterial phyla were differentially abundant across planting time at each site. At the Kansas site, Firmicutes (−3.95 log2 Fold Change (lfc)), Bacteroidota (4.43 lfc) and Actinobacteriota (2.79 lfc) were differentially abundant in regular vs. early planting, while only Actinobacteriota (1.74 lfc) was also differentially abundant in regular vs. early planting at the South Dakota site (Fig. 1c, Fig. S6). Fungal phyla Basidiomycota and Ascomycota were differentially abundant across planting time, but the direction of change was different between Kansas and South Dakota. Ascomycota abundance was higher in early planting in South Dakota and lower in Kansas, while Basidiomycota showed the opposite pattern (Fig 1d, Fig. S7). A total of 93 fungal genera were differentially abundant in Kansas samples, 72 more abundant in early planted and 21 in regular planted sorghum. For the South Dakota field samples, 8 fungal genera were significantly higher in abundance in early planted samples and 57 in regular planted samples, for a total of 65 significantly differentially abundant fungi genera (Fig. S7). Fungal genera with consistent differential abundance patterns included *Mycosphaerella* with higher abundance in early compared to regular planting and *Capnocheirides, Spegazzinia, Pseudozyma, Ustilago, Pseudopithiomyces, Cladosporium* with higher abundance in regular compared to early planting samples.

### Sorghum accessions tested had varying levels of microbiome-dependent chilling tolerance

A selected set of eight accessions (Tables 1 and 2) and the chilling sensitive check RTx430 were evaluated for chilling tolerance in the presence/absence of microbiomes in in a controlled environment growth chamber experiment. The goal here was to evaluate if microbiomes contributed at least in part to chilling tolerance phenotypes exhibited by these accessions. Chilling stress significantly reduced all plant growth traits evaluated irrespective of the presence or absence of microbiomes and the extent of reduction varied significantly between accessions (Table S4). Chlorophyll index was not significantly affected by the microbiome treatment in both control and chilling conditions (Table S4). However, accessions differed significantly within temperature treatments, with significant interaction between accession and microbiome treatment was observed only under control conditions (P<0.001) (Table S4). Quantum Yield was significantly affected by the microbiome (P<0.01-0.05), accession (P<0.001), and their interaction (P<0.001) both under control and chilling conditions (Table S4).

Leaf area was significantly affected by microbiome (P<0.05), accession (P<0.01) and their interactions (P<0.001) under both control and chilling conditions (Table S4). In the microbiome inoculation treatment, PI566819, PI642998, PI656019, and PI656121 had significantly (P<0.05) higher leaf area under the control condition (Fig. 2A). In contrast, the microbiome inoculation reduced (P<0.05) the leaf area under chilling stress in PI597971, PI642998, and PI656019 (Fig. 2B). Assessment based on microbiome response index suggested that microbiome inoculation had a positive impact on leaf area both under control and chilling stress conditions for accessions PI534117, PI656014, PI656121 (Fig. 2C). The microbiome inoculation promoted leaf area under control conditions but reduced leaf area under chilling stress conditions in PI566819, PI597971, PI642998, PI656019 and RTx430 (Fig. 2C). In one accession, PI576376, the microbiome inoculation reduced the leaf area under both control and chilling stress (Fig. 2C).

Similarly, seedling shoot biomass was significantly affected by inoculation treatment (P<0.05), accession P<0.01) and their interactions (P<0.001) under control conditions. In the chilling condition, seedling biomass was only affected by accession (Table S4). On average, shoot biomass across accessions with microbiome inoculation was increased by 53% in control and 17.5% in chilling stress conditions (Fig. 2D, E). The microbiome inoculation promoted shoot biomass under control conditions, but reduced biomass under chilling stress conditions in PI534117, PI566819, PI597971, PI642998 and PI656019 (Fig. 2F). Whereas the microbiome inoculation had a positive impact on biomass both under control and chilling stress conditions in PI656014, PI656121 and RTx430 (Fig. 2F). On the other hand, microbial inoculation had a positive effect on leaf area and shoot biomass under both control and chilling stress conditions in PI656014 and PI656121 (Fig. 2C, F). The microbiome had a largely neutral or negative effect on chlorophyll content and Quantum Yield (Fig. S3).

CTIM minus CTIm values indicate the level of microbiome-dependent chilling response for each of the measured traits (Fig. 3). A positive microbiome-dependent chilling tolerance was observed for chlorophyll index in three accessions PI576376, PI642998, PI656014. Three accessions, PI534117, PI656014 and PI656121, displayed enhanced microbiome-dependent chilling tolerance for leaf area. For shoot biomass, PI576376, PI656014 PI656121, and RTx430 showed microbiome-dependent chilling tolerance (Fig. 3). No positive responses were observed for QY, although PI576376, PI656014, and PI642998 had the least negative response to microbial inoculation (Fig. 3). Apart from QY, one accession (PI656014) showed positive microbiome-dependent chilling tolerance for all other traits. All other accessions that showed microbiome-dependent chilling tolerance rank as follows: PI656121 positive index for leaf area, shoot biomass; PI576376 positive index for shoot biomass, chlorophyll; RTx430 positive index for shoot biomass; PI642998 positive index for chlorophyll; PI534117 positive index for leaf area.

**Figure 3.**
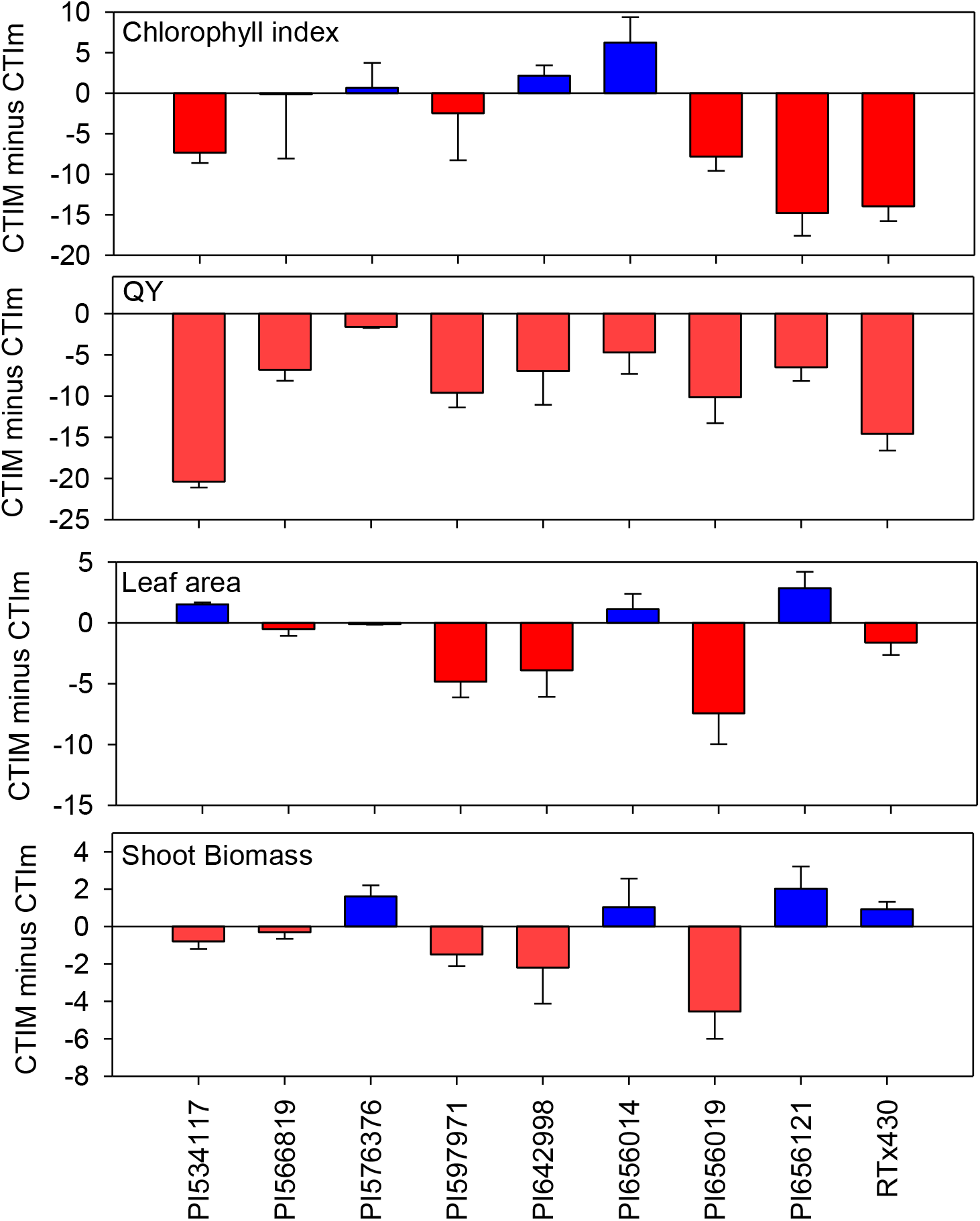
CTIM minus CTIm values for chlorophyll index, QY, leaf area and shoot biomass for nine sorghum accessions evaluated under growth chamber conditions. Accessions with blue are the ones showing microbiome-dependent chilling tolerance for traits and red indicates *vice versa*. CTIm - Chilling Tolerance Index with no microbe inoculum; CTIM - Chilling Tolerance Index with the microbe inoculum.

## Discussion

Early season chilling stress is a yield-limiting factor for sorghum in temperate climates. Tapping into sorghum genetic resources and microbes that enhance chilling tolerance could expand solutions for such yield limitations. This study identified sorghum accessions with microbiome-dependent responses to chilling stress. Overall, sorghum response to temperature treatments was accession-specific in both field and growth chamber experiments. In the field, early planting imposed significant chilling stress on sorghum plants, resulting in an overall 70-90% decrease in shoot biomass and leaf area compared to regular planting. A similar trend was found for sorghum in the growth chamber and field-based temperature treatments in previous studies (Chiluwal *et al*. 2018, Moghimi *et al*. 2019, Ostmeyer *et al*. 2020).

The performance of sorghum accessions was broadly similar across the field sites. Generally, accessions with higher shoot biomass, leaf area, and percent emergence in early planting in Kansas also tended to have higher trait values in South Dakota. Sorghum accessions that had the highest leaf area and shoot biomass in early planting conditions also tended to perform better in growth chamber chilling stress in the absence of microbial inoculation (PI577356, PI642998). This suggests that microbial inoculations can have a negative growth impact on some accessions. Interestingly, these genotypes performed much worse in growth chamber chilling stress with microbial inoculation.

Phyllosphere microbial communities associated with chilling-stressed sorghum were significantly different from microbes on optimally grown sorghum leaves in the field experiment. The microbial community differences between early and regular planting could be due to seasonality, temperature, or differences in plant stress among many other environmental factors. These data provide an important baseline for further work on chilling stress and plant-microbiome interactions. Understanding mechanisms for traits like microbe-enhanced chilling stress tolerance requires greenhouse and lab conditions, which often result in plant microbiome dynamics that are quite different from field conditions. Comparison of the field plant microbiome and greenhouse plant microbiome in future studies may pinpoint important taxa or community dynamics.

Currently there are few studies on the role of phyllosphere microbes in plant chilling stress tolerance. Two other studies have characterized sorghum phyllosphere microbial communities, but these focused on drought stress (Gao *et al*. 2020, Sun *et al*. 2021). For comparison, we identified similar sorghum phyllosphere taxa, such as *Alternaria* and *Cladosporium* dominating fungal communities (Gao *et al*. 2020) and Proteobacteria and Actinobacteria dominating bacterial communties (Sun *et al*. 2021). One other study has examined the effect of chilling stress on a plant microbiome. Beirinckx *et al*. (2020) characterized bacterial endophytes in maize roots enriched by chilling stress in pot experiments. Although this maize study examined roots in pots and we examined leaf microbiomes in the field, we found similar groups of bacterial taxa were more abundant in early planting (chilling conditions) as their chilling conditions in a pot experiment. These included bacterial families Caulobacteraceae and Oxalobacteraceae identified as more abundant in early planting (See Table S5). We also found that Actinobacteria were more abundant in regular planting compared to early planting leaf samples at both field sites, which reflects the finding that Actinobacteria are depleted under chilling conditions in pot experiments (Beirinckx *et al*. 2020). The abundant and differentially abundant taxa included several characterized fungal endophytes. For example, *Epicoccum dendrobii* is an endophyte with potential for biological control against a fungal plant pathogen (Bian *et al*. 2021) and was prevalent at regular planting in Kansas. We found that *Vishniacozyma globispora*, a basidiomycete yeast, was more abundant in early planting sorghum leaves. *Vishniacozyma* species have been identified as rice endophytes, Arctic plant endophytes, and tropical forest tree endophytes (Tantirungkij *et al*. 2015, Santiago *et al*. 2017, Felix *et al*. 2020). Another close relative is a psychrotolerant yeast isolated from a glacier (Tsuji *et al*. 2019). Basidomycete yeasts may be potentially important fungal endophytes for improving plant stress tolerance.

Plants are thought to primarily obtain their microbiomes from the soil (Beirinckx *et al*. 2020). The microbial vs. control inoculation treatments in the growth chamber experiment likely created a large difference in soil microbial diversity available to the plant. The sterilized potting media should harbor very few bacteria and fungi, while soil-extracted inoculum added diverse soil microbial taxa to the potting media. The temperature has been shown to shift bacterial and fungal community composition in soils and plants (Beirinckx *et al*. 2020, Cloutier *et al*. 2020), and we would assume such differences occurred in the field and growth chamber plant microbiomes. In this study, we focus on the plant physiological outcome of these microbial treatments to assess the potential for microbial-enhanced chilling tolerance in sorghum.

PI656014 and PI656121 showed a positive index for microbiome-dependent chilling tolerance in the growth chamber (Fig. 4) and these accessions were neither the best nor worst-performing accessions in the field. For example, PI576376 had the highest shoot biomass in early planting in Kansas and the fourth highest in South Dakota, and showed a positive shoot biomass response to microbial inoculation in chilling stress. The accessions with greater shoot row biomass in early planting field conditions (PI656014, PI576376) also had positive microbiome-dependent chilling trait responses in the growth chamber experiment (PI576376: biomass, chlorophyll, PI656014: biomass, chlorophyll, leaf area). This suggests that some accessions will be amenable to chilling tolerance enhancements via microbial inoculants.

To our knowledge, this is the first study to compare microbial inoculation treatments across plant genotypes under early-stage chilling stress. We observed diverse responses among sorghum accessions to microbial inoculation that were temperature condition dependent. For example, most accessions had increased leaf area and shoot biomass with microbial inoculation under control (30°C/20°C, day/night) temperatures, with 3 and 4 accessions exhibiting a statistically significant response in leaf area and shoot biomass, respectively (Fig. 3A, D). Under chilling stress (20°C/15°C, day/night), a positive effect of microbial inoculation on at least one trait is observed qualitatively in six accessions out of nine. PI656014 performed best under chilling stress: three out of four traits had higher values with microbial inoculation. Other accessions showed different combinations of traits that respond positively to microbe inoculation under chilling stress. Overall, the diverse trait response of accessions to microbial inoculation in the growth chamber under different temperature conditions suggests that microbes may enhance plant chilling tolerance via genotype-specific mechanisms.

Although few other studies use a diverse microbiome, several studies have tested the effect of individual microbial taxa on plants under chilling stress. These studies show similar plant biomass effects to our study, where the plant growth-promoting bacteria increase the biomass of the inoculated plant relative to non-inoculated plant under chilling stress, while the microbe-inoculated plants in the control temperatures result in an even greater plant biomass increase relative to non-inoculated plants (Barka *et al*. 2006, Subramanian *et al*. 2016, Wang *et al*. 2016). Therefore, further exploration of individual or consortia of microbes and genetic loci responsible for their microbial recruitment and benefit have great promise to enhance chilling tolerance in sorghum.

## Conclusion

Pinning down microbial effects on plant phenotype can be difficult due to context-dependent responses and variation in experimental microbial communities. In spite of this variability, we observe accession-specific positive and negative responses to high and low microbial diversity treatments under chilling stress temperatures, which we have termed the microbiome-dependent chilling stress response. Follow-up studies will screen microbial taxa with the diverse panel of sorghum accessions for microbe taxa impact on accession chilling tolerance in the growth chamber and field experiments. Candidate taxa and sorghum accessions will be assessed for the genetic basis of microbiome-dependent chilling tolerance.

## Acknowledgements

Contribution number 21-259-J from Kansas Agricultural Experiment Station. This work was also supported by the USDA National Institute of Food and Agriculture, hatch multistate project 1014561, grant awards from the National Science Foundation/EPSCoR Cooperative Agreement #1849206; National Science Foundation’s Plant Genome Research Program (IOS-1350189); and SD Agricultural Experiment Station (SD00H697-20).

## References

Abarenkov, Kessy; Zirk, Allan; Piirmann, Timo; Pöhönen, Raivo; Ivanov, Filipp; Nilsson, R. Henrik; Kõljalg, Urmas (2020): UNITE general FASTA release for Fungi. Version 04.02.2020. UNITE Community. https://doi.org/10.15156/BIO/786368.

Backer, R., Rokem, J. S., Ilangumaran, G., Lamont, J., Praslickova, D., Ricci, E., … Smith, D. L. (2018). Plant Growth-Promoting Rhizobacteria: Context, Mechanisms of Action, and Roadmap to Commercialization of Biostimulants for Sustainable Agriculture. Frontiers in Plant Science, 9. doi: 10.3389/fpls.2018.01473

Barka, E. A., Nowak, J., & Clément, C. (2006). Enhancement of Chilling Resistance of Inoculated Grapevine Plantlets with a Plant Growth-Promoting Rhizobacterium, Burkholderia phytofirmans Strain PsJN. Applied and Environmental Microbiology, 72(11), 7246–7252. doi: 10.1128/AEM.01047-06

Beirinckx, S., Viaene, T., Haegeman, A., Debode, J., Amery, F., Vandenabeele, S., … Goormachtig, S. (2020). Tapping into the maize root microbiome to identify bacteria that promote growth under chilling conditions. Microbiome, 8(1), 54. doi: 10.1186/s40168-020-00833-w

Bian, J.-Y., Fang, Y.-L., Song, Q., Sun, M.-L., Yang, J.-Y., Ju, Y.-W., … Huang, L. (2020). The Fungal Endophyte Epicoccum dendrobii as a Potential Biocontrol Agent Against Colletotrichum gloeosporioides. Phytopathology®, 111(2), 293–303. doi: 10.1094/PHYTO-05-20-0170-R

Callahan, B. J., McMurdie, P. J., Rosen, M. J., Han, A. W., Johnson, A. J. A., & Holmes, S. P. (2016). DADA2: High-resolution sample inference from Illumina amplicon data. Nature Methods, 13(7), 581–583. doi: 10.1038/nmeth.3869

Chelius, M. K., & Triplett, E. W. (2001). The Diversity of Archaea and Bacteria in Association with the Roots of Zea mays L. Microbial Ecology, 41(3), 252–263. doi: 10.1007/s002480000087

Chiluwal, A., Bheemanahalli, R., Perumal, R., Asebedo, A. R., Bashir, E., Lamsal, A., … Krishna Jagadish, S. V. (2018). Integrated aerial and destructive phenotyping differentiates chilling stress tolerance during early seedling growth in sorghum. Field Crops Research, 227, 1–10. doi: 10.1016/j.fcr.2018.07.011

Chopra, R., Burow, G., Hayes, C., Emendack, Y., Xin, Z., & Burke, J. (2015). Transcriptome profiling and validation of gene based single nucleotide polymorphisms (SNPs) in sorghum genotypes with contrasting responses to cold stress. BMC Genomics, 16(1), 1040. doi: 10.1186/s12864-015-2268-8

Cloutier, M., Chatterjee, D., Elango, D., Cui, J., Bruns, M. A., & Chopra, S. (2020). Sorghum Root Flavonoid Chemistry, Cultivar, and Frost Stress Effects on Rhizosphere Bacteria and Fungi. Phytobiomes Journal, PBIOMES-01-20-0013-FI. doi: 10.1094/PBIOMES-01-20-0013-FI

Ding, S., Huang, C.-L., Sheng, H.-M., Song, C.-L., Li, Y.-B., & An, L.-Z. (2011). Effect of inoculation with the endophyte Clavibacter sp. Strain Enf12 on chilling tolerance in Chorispora bungeana. Physiologia Plantarum, 141(2), 141–151. doi: https://doi.org/10.1111/j.1399-3054.2010.01428.x

Gao, C., Montoya, L., Xu, L., Madera, M., Hollingsworth, J., Purdom, E., … Taylor, J. W. (2020). Fungal community assembly in drought-stressed sorghum shows stochasticity, selection, and universal ecological dynamics. Nature Communications, 11(1), 1–14. doi: 10.1038/s41467-019-13913-9

Gardes, M., & Bruns, T. D. (1993). ITS primers with enhanced specificity for basidiomycetes— Application to the identification of mycorrhizae and rusts. Molecular Ecology, 2(2), 113–118. doi: 10.1111/j.1365-294X.1993.tb00005.x

Kang, S.-M., Khan, A. L., Waqas, M., You, Y.-H., Hamayun, M., Joo, G.-J., … Lee, I.-J. (2015). Gibberellin-producing Serratia nematodiphila PEJ1011 ameliorates low temperature stress in Capsicum annuum L. European Journal of Soil Biology, 68, 85–93. doi: 10.1016/j.ejsobi.2015.02.005

Lata, R., Chowdhury, S., Gond, S. K., & White, J. F. (2018). Induction of abiotic stress tolerance in plants by endophytic microbes. Letters in Applied Microbiology, 66(4), 268–276. doi: https://doi.org/10.1111/lam.12855

Lundberg, D. S., Yourstone, S., Mieczkowski, P., Jones, C. D., & Dangl, J. L. (2013). Practical innovations for high-throughput amplicon sequencing. Nature Methods, 10(10), 999–1002. doi: 10.1038/nmeth.2634

Marla, S. R., Burow, G., Chopra, R., Hayes, C., Olatoye, M. O., Felderhoff, T., … Morris, G. P. (2019). Genetic Architecture of Chilling Tolerance in Sorghum Dissected with a Nested Association Mapping Population. G3 Genes|Genomes||Genetics, 9(12), 4045–4057. doi: 10.1534/g3.119.400353

Martin, M. (2011). Cutadapt removes adapter sequences from high-throughput sequencing reads. EMBnet.Journal, 17(1), 10–12. doi: 10.14806/ej.17.1.200

Maulana, F., & Tesso, T. T. (2013). Cold Temperature Episode at Seedling and Flowering Stages Reduces Growth and Yield Components in Sorghum. Crop Science, 53(2), 564–574. doi: https://doi.org/10.2135/cropsci2011.12.0649

Moghimi, N., Desai, J. S., Bheemanahalli, R., Impa, S. M., Vennapusa, A. R., Sebela, D., … Jagadish, S. V. K. (2019). New candidate loci and marker genes on chromosome 7 for improved chilling tolerance in sorghum. Journal of Experimental Botany, 70(12), 3357–3371. doi: 10.1093/jxb/erz143

Ostmeyer, T., Bheemanahalli, R., Srikanthan, D., Bean, S., Peiris, K. H. S., Madasamy, P., … Jagadish, S. V. K. (2020). Quantifying the agronomic performance of new grain sorghum hybrids for enhanced early-stage chilling tolerance. Field Crops Research, 258, 107955. doi: 10.1016/j.fcr.2020.107955

Raymundo, R., Ciampitti, I. A., & Morris, G. (2021). Crop modeling defines opportunities and challenges for drought escape, water capture, and yield increase using chilling-tolerant sorghum. BioRxiv, 2021.01.27.428532. doi: 10.1101/2021.01.27.428532

South Dakota Mesonet, South Dakota State University (2021). SD Mesonet Archive. Retrieved from https://mesonet.sdstate.edu/archive. (n.d.).

Subramanian, P., Kim, K., Krishnamoorthy, R., Mageswari, A., Selvakumar, G., & Sa, T. (2016). Cold Stress Tolerance in Psychrotolerant Soil Bacteria and Their Conferred Chilling Resistance in Tomato (Solanum lycopersicum Mill.) under Low Temperatures. PLOS ONE, 11(8), e0161592. doi: 10.1371/journal.pone.0161592

Sun, A., Jiao, X.-Y., Chen, Q., Wu, A.-L., Zheng, Y., Lin, Y.-X., … Hu, H.-W. (2021). Microbial communities in crop phyllosphere and root endosphere are more resistant than soil microbiota to fertilization. Soil Biology and Biochemistry, 153, 108113. doi: 10.1016/j.soilbio.2020.108113

Tantirungkij, M., Nasanit, R., & Limtong, S. (2015). Assessment of endophytic yeast diversity in rice leaves by a culture-independent approach. Antonie van Leeuwenhoek, 108(3), 633–647. doi: 10.1007/s10482-015-0519-y

Tari, I., Laskay, G., Takács, Z., & Poór, P. (2013). Response of Sorghum to Abiotic Stresses: A Review. Journal of Agronomy and Crop Science, 199(4), 264–274. doi: https://doi.org/10.1111/jac.12017

Wagner, M. R., Lundberg, D. S., del Rio, T. G., Tringe, S. G., Dangl, J. L., & Mitchell-Olds, T. (2016). Host genotype and age shape the leaf and root microbiomes of a wild perennial plant. Nature Communications, 7, 12151. doi: 10.1038/ncomms12151

Wallace, J. G., Kremling, K. A., Kovar, L. L., & Buckler, E. S. (2018). Quantitative Genetics of the Maize Leaf Microbiome. Phytobiomes Journal, 2(4), 208–224. doi: 10.1094/PBIOMES-02-18-0008-R

Wallace, J., Laforest-Lapointe, I., & Kembel, S. W. (2018). Variation in the leaf and root microbiome of sugar maple (Acer saccharum) at an elevational range limit. PeerJ, 6. doi: 10.7717/peerj.5293

Wang, C., Wang, C., Gao, Y.-L., Wang, Y.-P., & Guo, J.-H. (2016). A Consortium of Three Plant Growth-Promoting Rhizobacterium Strains Acclimates Lycopersicon esculentum and Confers a Better Tolerance to Chilling Stress. Journal of Plant Growth Regulation, 35(1), 54–64. doi: 10.1007/s00344-015-9506-9

White, T. J., Bruns, T., Lee, S., & Taylor, J. (1990). Amplification and direct sequencing of ribosomal RNA genes for phylogenetics. In M. A. Innis, D. H. Gelfand, J. J. Sninsky, & T. J. White (Eds.), PCR Protocols (pp. 315–322). San Diego: Academic Press. Retrieved from http://www.sciencedirect.com/science/article/pii/B9780123721808500421

Yu, J., & Tuinstra, M. R. (2001). Genetic Analysis of Seedling Growth under Cold Temperature Stress in Grain Sorghum. Crop Science, 41(5), 1438–1443. doi: https://doi.org/10.2135/cropsci2001.4151438x

Zhou, L., Li, C., White, J. F., & Johnson, R. D. (2021). Synergism between calcium nitrate applications and fungal endophytes to increase sugar concentration in Festuca sinensis under cold stress. PeerJ, 9, e10568. doi: 10.7717/peerj.10568

