## Supplementary information for "Evidence for microbiome-dependent chilling tolerance in sorghum"

### Supplementary Figures

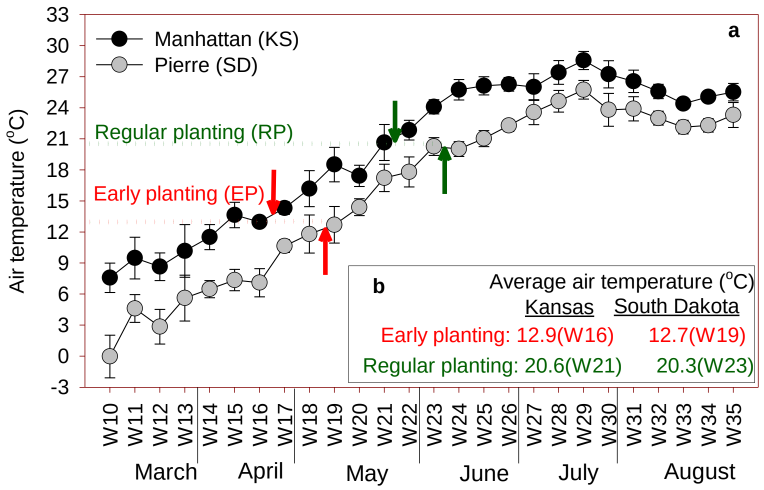


**Supplementary Figure 1**. (a) Observed weekly average air temperatures (2010-2018) for Julian weeks 10 to 35 at Manhattan, KS and Pierre, SD, the locations to be used for this research. Arrows indicate proposed planting dates for each location based on (b) average temperatures of 12.8°C for Early Planting (EP) and 20.3 °C Regular/Optimum Planting (RP).

(A)

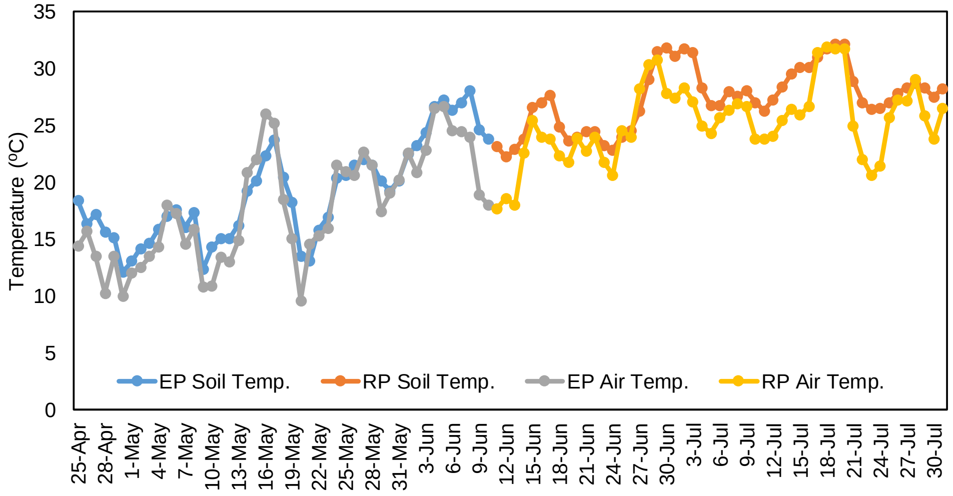


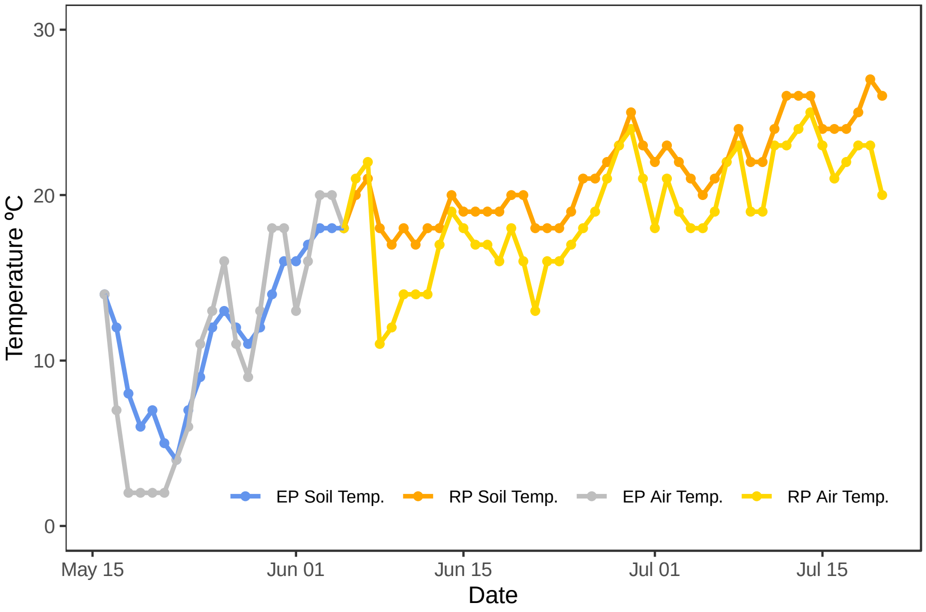


(B)

**Supplementary Figure 2**. Average daily air and soil temperature in the field under early and regular planting conditions at the Kansas (A) and South Dakota (B) field sites.

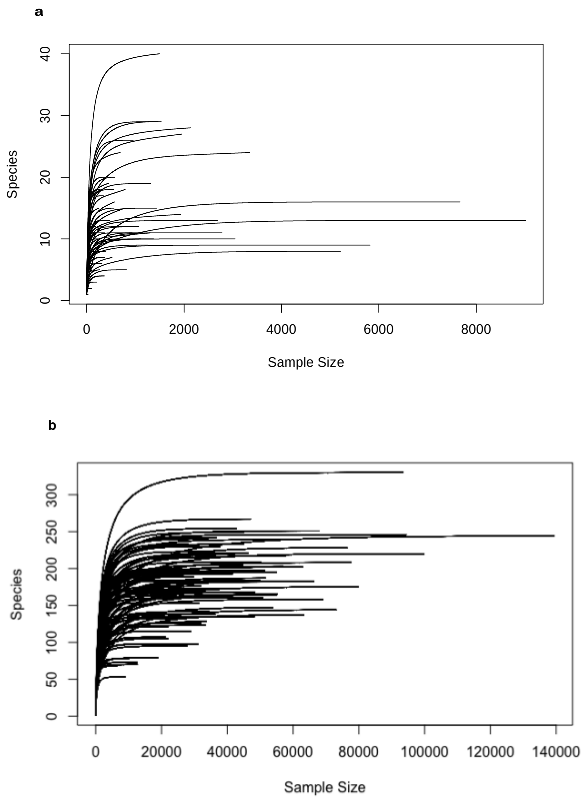


**Supplementary Figure 3.** Rarefaction curves for bacterial (a) and fungal (b) amplicon sequencing libraries show that the sequencing depth of most libraries likely detected the total number of taxa in each leaf sample.

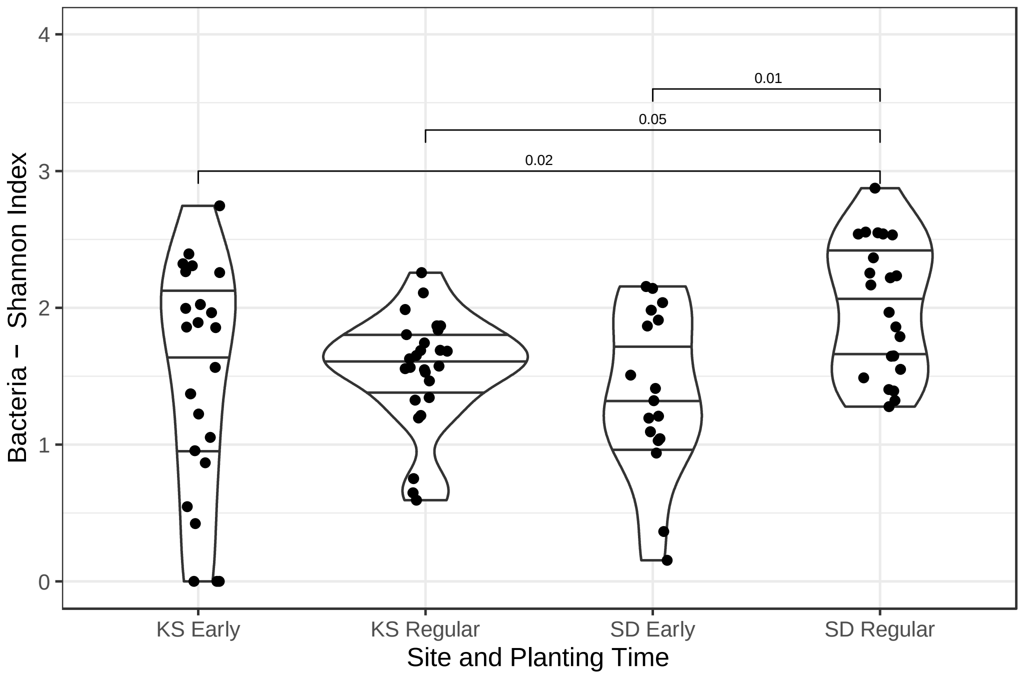


**Supplementary Figure 4.** Violin plot shows bacteria sample Shannon index by site and planting time with results from ANOVA and Tukey's HSD. Within the Kansas site, diversity is the same between early and regular planting time. Within South Dakota samples, diversity is higher in regular planting samples than early. Lines within violins are quartiles and points are samples.

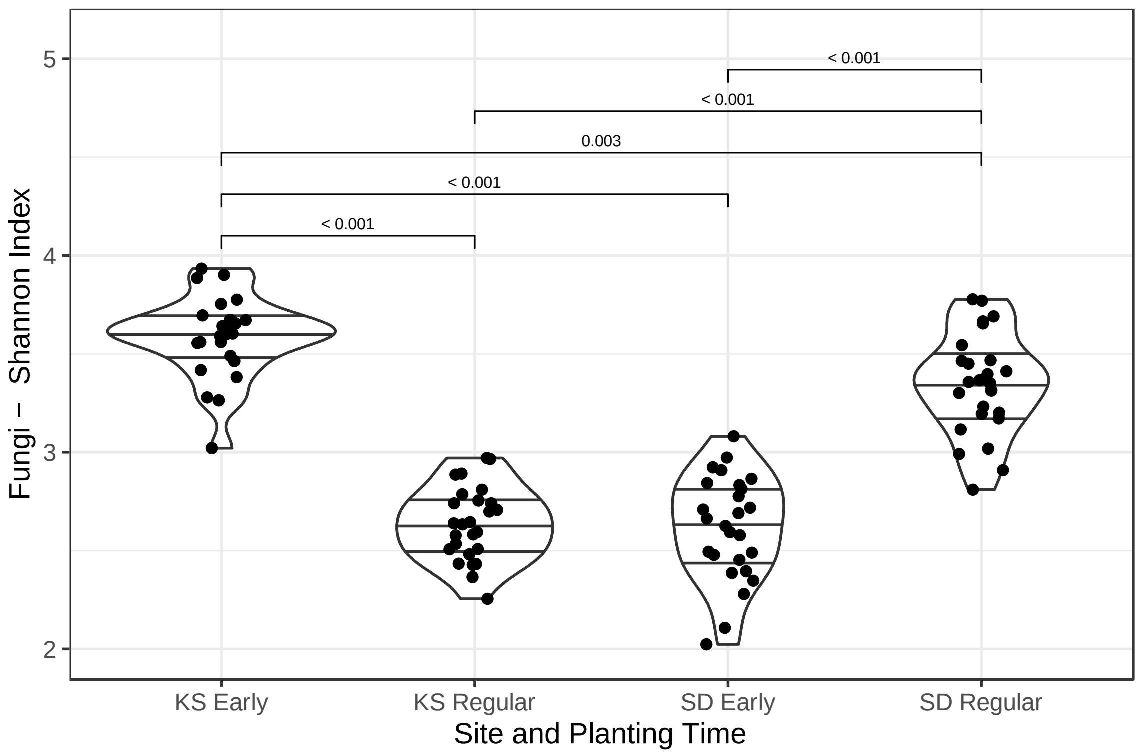


**Supplementary Figure 5.** Violin plot shows fungi sample Shannon index by site and planting time with results from ANOVA and Tukey’s HSD. Within the Kansas site, diversity is higher at early planting than regular planting time. Within South Dakota samples, diversity is higher in regular planting samples than early. Lines within violins are quartiles and points are samples.

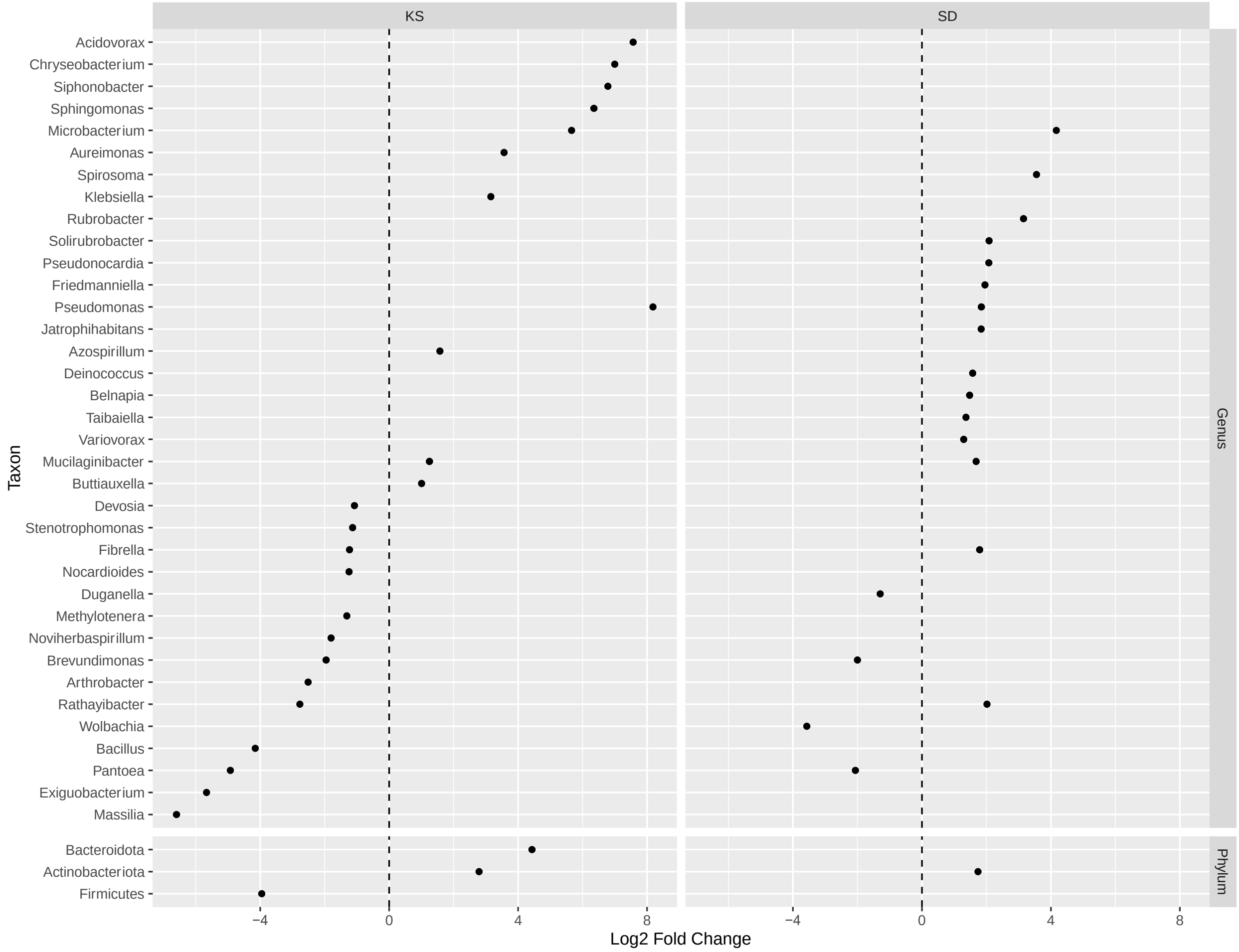


**Supplemental Figure 6**. Log2 Fold Change of bacterial genera and phyla significantly differentially abundant (padj <0.05) between early planting (negative lfc) and regular planting (positive lfc) at each field site.

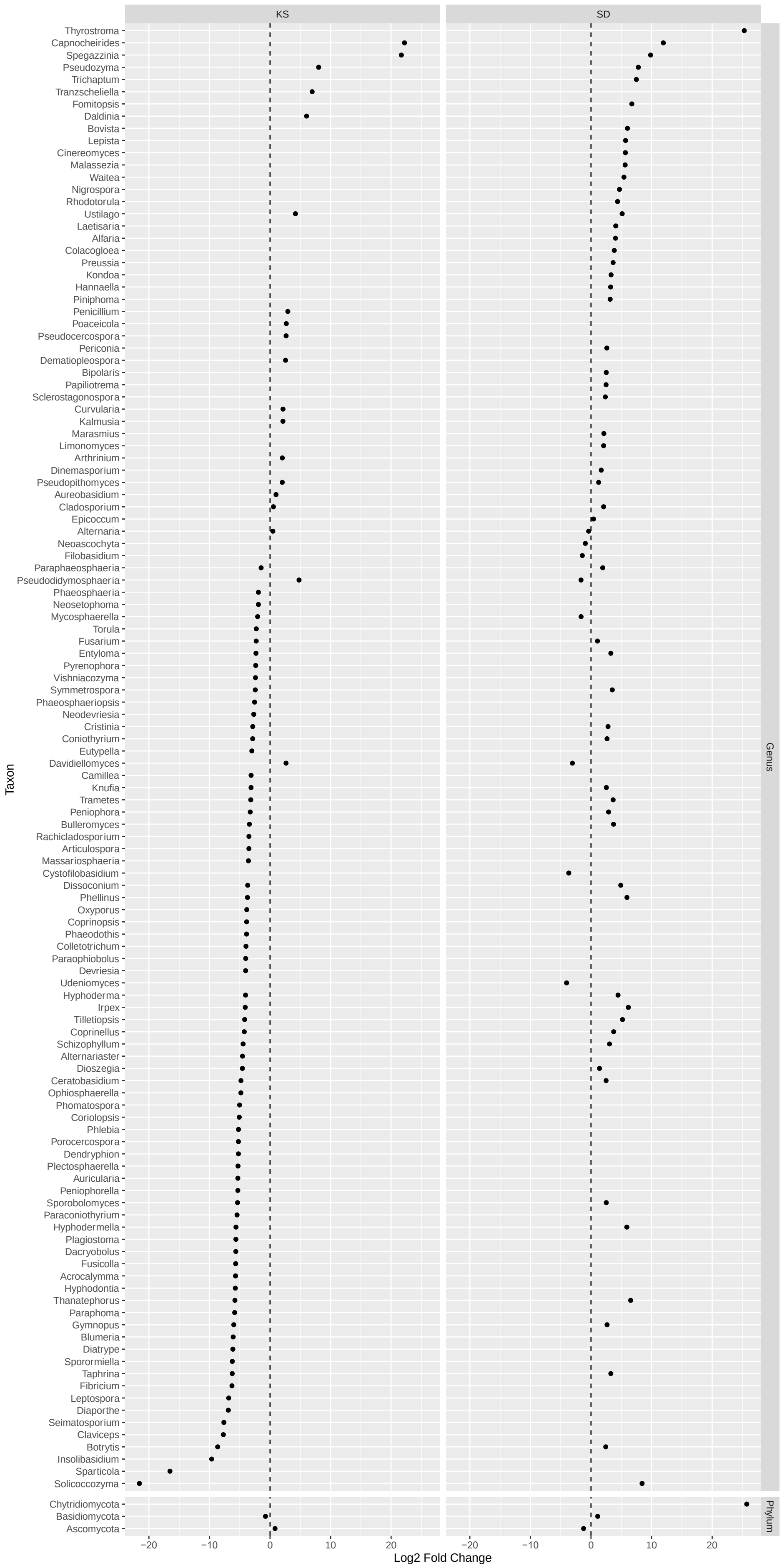

KS
SD
Genus
Phylum
Taxon
Log2 Fold Change
−20
−10
0
10
20
Thyrostroma
Capnocheirides
Spegazzinia
Pseudozyma
Trichaptum
Tranzscheliella
Fomitopsis
Daldinia
Bovista
Lepista
Cinereomyces
Malassezia
Waitea
Nigrospora
Rhodotorula
Ustilago
Laetisaria
Alfaria
Colacogloea
Preussia
Kondoa
Hannaella
Piniphoma
Penicillium
Poaceicola
Pseudocercospora
Periconia
Dematiopleospora
Bipolaris
Papiliotrema
Sclerostagonospora
Curvularia
Kalmusia
Marasmius
Limonomyces
Arthrinium
Dinemasporium
Pseudopithomyces
Aureobasidium
Cladosporium
Epicoccum
Alternaria
Neoascochyta
Filobasidium
Paraphaeosphaeria
Pseudodidymosphaeria
Phaeosphaeria
Neosetophoma
Mycosphaerella
Torula
Fusarium
Entyloma
Pyrenophora
Vishniacozyma
Symmetrospora
Phaeosphaeriopsis
Neodevriesia
Cristinia
Coniothyrium
Eutypella
Davidiellomyces
Camillea
Knufia
Trametes
Peniophora
Bulleromyces
Rachicladosporium
Articulospora
Massariosphaeria
Cystofilobasidium
Dissoconium
Phellinus
Oxyporus
Coprinopsis
Phaeodothis
Colletotrichum
Paraophiobolus
Devriesia
Udeniomyces
Hyphoderma
Irpex
Tilletiopsis
Coprinellus
Schizophyllum
Alternariaster
Dioszegia
Ceratobasidium
Ophiosphaerella
Phomatospora
Coriolopsis
Phlebia
Porocercospora
Dendryphion
Plectosphaerella
Auricularia
Peniophorella
Sporobolomyces
Paraconiothyrium
Hyphodermella
Plagiostoma
Dacryobolus
Fusicolla
Acrocalymma
Hyphodontia
Thanatephorus
Paraphoma
Gymnopus
Blumeria
Diatrype
Sporormiella
Taphrina
Fibricium
Leptospora
Diaporthe
Seimatosporium
Claviceps
Botrytis
Insolibasidium
Sparticola
Solicoccozyma
Chytridiomycota
Basidiomycota
Ascomycota

**Supplemental Figure 7**. Log2 Fold Change of fungal genera and phyla significantly differentially abundant (padj <0.05) between early planting (negative lfc) and regular planting (positive lfc) at each field site.

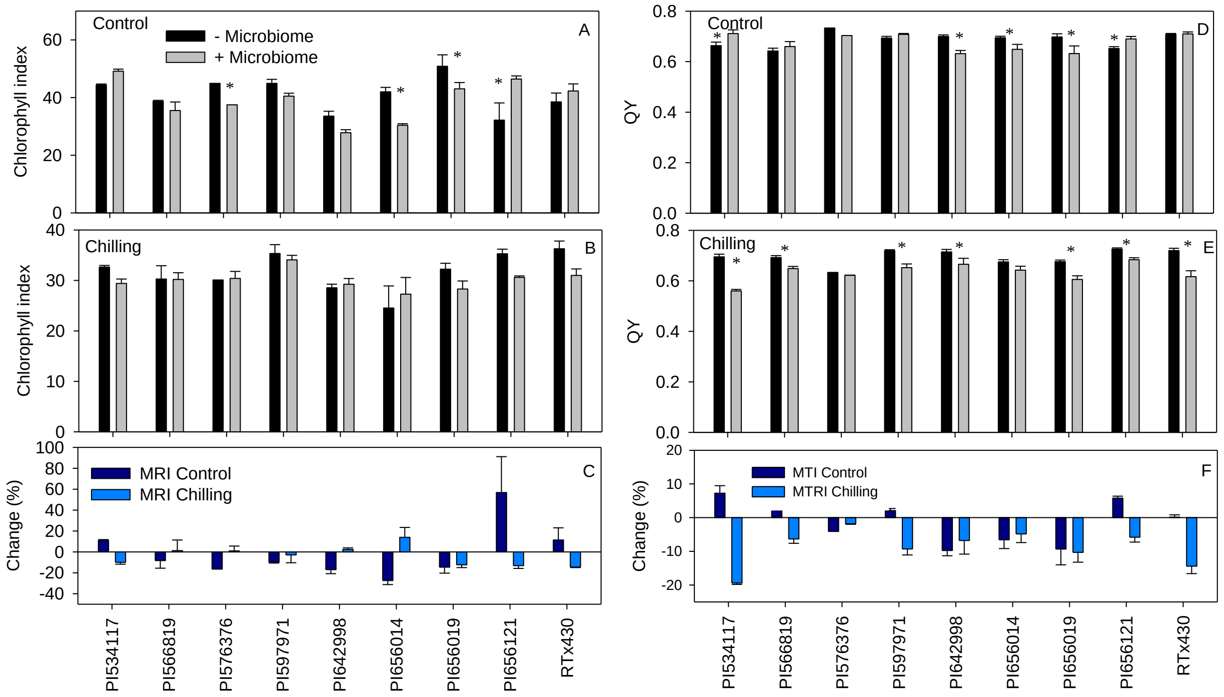


**Supplementary Figure 8.** Chlorophyll index (A-C) and quantum yield (D-F) of sorghum accessions at 45 DAP with and without microbial inoculation under control (A and D; 30ºC/20ºC, day/night) and chilling stress conditions (B and E; 20ºC/10ºC, day/night). Microbiome Response Index (MRI) indicates the percentage change with and without microbial inoculation under control and chilling conditions (C and F). * represents a significant difference between treatments (microbiome and accession) at 5% LSD.

### Supplementary Tables

**Supplementary Table 1**. Relative abundance of ten most abundant bacterial genera by planting time and site.

**Bacteria**

| Kansas | | | | South Dakota | | | |
|---|---|---|---|---|---|---|---|
| **Genus** | **Early Planting** | **Genus** | **Regular Planting** | **Genus** | **Early Planting** | **Genus** | **Regular Planting** |
| *Exiguobacterium* | 0.301 | *Pseudomonas* | 0.259 | *Pantoea* | 0.420 | *Microbacterium* | 0.205 |
| *Massilia* | 0.244 | *Microbacterium* | 0.228 | *Buchnera* | 0.103 | *Rathayibacter* | 0.159 |
| *Paenibacillus* | 0.132 | *Acidovorax* | 0.201 | *Hymenobacter* | 0.100 | *Spirosoma* | 0.100 |
| *Domibacillus* | 0.088 | *Sphingomonas* | 0.104 | *Massilia* | 0.065 | *Pantoea* | 0.085 |
| *Pantoea* | 0.049 | *Chryseobacterium* | 0.085 | *Acinetobacter* | 0.057 | *Pseudomonas* | 0.076 |
| *Bacillus* | 0.048 | *Siphonobacter* | 0.074 | *Rathayibacter* | 0.042 | *Massilia* | 0.047 |
| *Buchnera* | 0.034 | *Allorhizobium-Neorhizobium-Pararhizobium-Rhizobium* | 0.013 | *Wolbachia* | 0.041 | *Pseudonocardia* | 0.036 |
| *Allorhizobium-Neorhizobium-Pararhizobium-Rhizobium* | 0.014 | *Aureimonas* | 0.007 | *Pseudomonas* | 0.022 | *Hymenobacter* | 0.035 |
| *Arthrobacter* | 0.012 | *Pedobacter* | 0.006 | *Brevundimonas* | 0.020 | *Rubrobacter* | 0.027 |
| *Rathayibacter* | 0.010 | *Klebsiella* | 0.005 | *Curtobacterium* | 0.020 | *Cutibacterium* | 0.022 |

**Supplementary Table 2**. Relative abundance of ten most abundant fungal genera by planting time and site.

**Fungi**

| Kansas | | | | South Dakota | | | |
|---|---|---|---|---|---|---|---|
| **Genus** | **Early Planting** | **Genus** | **Regular Planting** | **Genus** | **Early Planting** | **Genus** | **Regular Planting** |
| *Alternaria* | 0.223 | *Alternaria* | 0.444 | *Mycosphaerella* | 0.390 | *Alternaria* | 0.174 |
| *Mycosphaerella* | 0.133 | *Epicoccum* | 0.136 | *Alternaria* | 0.218 | *Epicoccum* | 0.158 |
| *Leptospora* | 0.130 | *Cladosporium* | 0.110 | *Epicoccum* | 0.118 | *Mycosphaerella* | 0.148 |
| *Epicoccum* | 0.095 | *Mycosphaerella* | 0.045 | *Neoascochyta* | 0.078 | *Ustilago* | 0.134 |
| *Sporobolomyces* | 0.068 | *Pseudopithomyces* | 0.040 | *Comoclathris* | 0.026 | *Neoascochyta* | 0.055 |
| *Cladosporium* | 0.050 | *Bipolaris* | 0.022 | *Filobasidium* | 0.024 | *Cladosporium* | 0.049 |
| *Neosetophoma* | 0.046 | *Curvularia* | 0.020 | *Vishniacozyma* | 0.013 | *Fusarium* | 0.020 |
| *Tilletiopsis* | 0.024 | *Neosetophoma* | 0.018 | *Cladosporium* | 0.013 | *Comoclathris* | 0.019 |
| *Filobasidium* | 0.017 | *Ustilago* | 0.016 | *Pseudodidymosphaeria* | 0.009 | *Ascochyta* | 0.018 |
| *Dioszegia* | 0.016 | *Neoascochyta* | 0.013 | *Fusarium* | 0.009 | *Filobasidium* | 0.015 |

**Supplementary Table 3**. Permutational ANOVA of bacterial and fungal communities by field site, planting time, and sorghum accession. Site, planting time and the site x planting time interaction significantly explained community dissimilarity.

| | Bacteria | | | | Fungi | | | |
|---|---|---|---|---|---|---|---|---|
| **Factor** | **df** | **F model** | **$R^2$** | **p** | **df** | **F model** | **$R^2$** | **p** |
| **Site** | 1 | 24.86 | 0.146 | *0.001* | 1 | 180.72 | 0.488 | *0.001* |
| **Planting time** | 1 | 33.70 | 0.197 | *0.001* | 1 | 60.528 | 0.164 | *0.001* |
| **Accession** | 8 | 1.42 | 0.066 | 0.084 | 8 | 1.088 | 0.024 | 0.354 |
| **Site x Planting time** | 1 | 21.00 | 0.123 | *0.001* | 1 | 29.662 | 0.08 | *0.001* |
| **Site x Accession** | 8 | 1.13 | 0.053 | 0.294 | 8 | 1.081 | 0.023 | 0.362 |
| **Planting time x Accession** | 8 | 1.22 | 0.057 | 0.208 | 8 | 0.911 | 0.02 | 0.576 |
| **Site x Planting time x Accession** | 7 | 1.14 | 0.047 | 0.264 | 8 | 0.94 | 0.02 | 0.55 |
| **Residuals** | 53 | | | | 67 | | | |
| **Total** | 87 | | | | 102 | | | |

**Supplementary Table 4.** Summary of analysis of variance for growth chambers experiment. The split-plot analysis was performed for each temperature treatment separately. Significant at the <0.05 (*), p<0.01 (**) and <0.001 (***) probability level; NS, Non-significant.

| | **Control** | **Chilling** |
|---|---|---|
| **Chlorophyll index** | | |
| Microbiome (M) | NS | NS |
| Accession (A) | <0.001 | <0.001 |
| MxA | <0.001 | NS |
| **QY** | | |
| Microbiome (M) | <0.05 | <0.01 |
| Accession (A) | <0.001 | <0.001 |
| MxA | <0.001 | <0.001 |
| **Leaf area ($cm^2$)** | | |
| Microbiome (M) | <0.05 | <0.05 |
| Accession (A) | <0.01 | <0.001 |
| MxA | <0.001 | <0.001 |
| **Shoot biomass (mg)** | | |
| Microbiome (M) | <0.01 | NS |
| Accession (A) | <0.001 | <0.01 |
| MxA | <0.05 | NS |

**Supplementary Table 5.** Differential abundance at the bacterial family level at each field site.

| | Kansas | | | | South Dakota | | |
|---|---|---|---|---|---|---|---|
| | Log2 Fold Change | LFC SE | padj | | Log2 Fold Change | LFC SE | padj |
| Oxalobacteraceae | -6.372 | 0.679 | 0.000 | Anaplasmataceae | -3.643 | 0.538 | 0.000 |
| Exiguobacteraceae | -5.662 | 0.527 | 0.000 | Erwiniaceae | -2.063 | 0.730 | 0.012 |
| Erwiniaceae | -4.886 | 0.706 | 0.000 | Caulobacteraceae | -2.011 | 0.522 | 0.001 |
| Bacillaceae | -4.111 | 0.515 | 0.000 | Intrasporangiaceae | -1.566 | 0.560 | 0.012 |
| Micrococcaceae | -3.679 | 0.603 | 0.000 | Propionibacteriaceae | 1.085 | 0.457 | 0.034 |
| Caulobacteraceae | -1.942 | 0.476 | 0.000 | Chitinophagaceae | 1.281 | 0.472 | 0.014 |
| Anaplasmataceae | -1.665 | 0.505 | 0.002 | Acetobacteraceae | 1.441 | 0.479 | 0.007 |
| Spirosomaceae | -1.370 | 0.583 | 0.031 | Deinococcaceae | 1.529 | 0.460 | 0.003 |
| Methylophilaceae | -1.332 | 0.403 | 0.002 | 67-14 | 1.534 | 0.480 | 0.004 |
| Intrasporangiaceae | -1.277 | 0.496 | 0.018 | Pseudomonadaceae | 1.773 | 0.671 | 0.017 |
| Nocardioidaceae | -1.243 | 0.430 | 0.008 | Frankiaceae | 1.802 | 0.483 | 0.001 |
| Xanthomonadaceae | -1.145 | 0.421 | 0.013 | Solirubrobacteraceae | 2.039 | 0.517 | 0.001 |
| Devosiaceae | -1.090 | 0.420 | 0.017 | Pseudonocardiaceae | 2.133 | 0.544 | 0.001 |
| Azospirillaceae | 1.528 | 0.444 | 0.002 | Microbacteriaceae | 2.341 | 0.705 | 0.003 |
| Rhizobiaceae | 1.739 | 0.617 | 0.010 | Comamonadaceae | 2.718 | 0.751 | 0.001 |
| Microbacteriaceae | 4.410 | 0.625 | 0.000 | Rubrobacteriaceae | 3.126 | 0.536 | 0.000 |
| Sphingomonadaceae | 6.254 | 0.474 | 0.000 | Spirosomaceae | 3.605 | 0.584 | 0.000 |
| Cytophagaceae | 6.680 | 0.621 | 0.000 | Micrococcaceae | 4.379 | 0.698 | 0.000 |
| Weeksellaceae | 6.907 | 0.553 | 0.000 | | | | |
| Comamonadaceae | 7.521 | 0.676 | 0.000 | | | | |
| Pseudomonadaceae | 8.127 | 0.636 | 0.000 | | | | |

**Supplementary Methods**

**Primer sequences** used in field sorghum leaf microbial community survey. All four versions of a primer were equimolarly pooled prior to use in PCR. The nucleotide frameshift was reported to improve base call quality by increasing nucleotide diversity in Lundberg *et al.* 2013. Frameshift nucleotides are in italics and locus specific sequences are in bold. The beginning of the primer sequence are Illumina technical sequences (adapters for the P5 and P7 primers in PCR2). 799F was based on Chelius & Triplett 2001, 1115R was based on Wallace *et al.* 2018a and Wallace *et al.* 2018b. ITS1F was based on Gardes & Bruns 1993, ITS2 was based on White *et al.* 1990.

| Primer name | Sequence |
|---|---|
| 799F_1 | TCGTCGGCAGCGTCAGATGTGTATAAGAGACAG**ACCMGGATTAGATACCCKG** |
| 799F_2 | TCGTCGGCAGCGTCAGATGTGTATAAGAGACAG*N***AACMGGATTAGATACCCKG** |
| 799F_3 | TCGTCGGCAGCGTCAGATGTGTATAAGAGACAG*NN***AACMGGATTAGATACCCKG** |
| 799F_4 | TCGTCGGCAGCGTCAGATGTGTATAAGAGACAG*NNN***AACMGGATTAGATACCCKG** |
| 1115R_1 | GTCTCGTGGGCTCGGAGATGTGTATAAGAGACAG**AGGGTTGCGCTCGTTG** |
| 1115R_2 | GTCTCGTGGGCTCGGAGATGTGTATAAGAGACAG*N***AGGGTTGCGCTCGTTG** |
| 1115R_3 | GTCTCGTGGGCTCGGAGATGTGTATAAGAGACAG*NN***AGGGTTGCGCTCGTTG** |
| 1115R_4 | GTCTCGTGGGCTCGGAGATGTGTATAAGAGACAG*NNN***AGGGTTGCGCTCGTTG** |
| ITS1F_1 | TCGTCGGCAGCGTCAGATGTGTATAAGAGACAG**CTTGGTCATTTAGAGGAAGTAA** |
| ITS1F_2 | TCGTCGGCAGCGTCAGATGTGTATAAGAGACAG*N***CTTGGTCATTTAGAGGAAGTAA** |
| ITS1F_3 | TCGTCGGCAGCGTCAGATGTGTATAAGAGACAG*NN***CTTGGTCATTTAGAGGAAGTAA** |
| ITS1F_4 | TCGTCGGCAGCGTCAGATGTGTATAAGAGACAG*NNN***CTTGGTCATTTAGAGGAAGTAA** |
| ITS2R_1 | GTCTCGTGGGCTCGGAGATGTGTATAAGAGACAG**GCTGCGTTCTTCATCGATGC** |
| ITS2R_2 | GTCTCGTGGGCTCGGAGATGTGTATAAGAGACAG*N***GCTGCGTTCTTCATCGATGC** |
| ITS2R_3 | GTCTCGTGGGCTCGGAGATGTGTATAAGAGACAG*NN***GCTGCGTTCTTCATCGATGC** |
| ITS2R_4 | GTCTCGTGGGCTCGGAGATGTGTATAAGAGACAG*NNN***GCTGCGTTCTTCATCGATGC** |


Chelius, M.K., and Triplett, E.W. (2001). The Diversity of Archaea and Bacteria in Association with the Roots of Zea mays L. Microb Ecol *41*, 252–263.

Gardes, M., and Bruns, T.D. (1993). ITS primers with enhanced specificity for basidiomycetes - application to the identification of mycorrhizae and rusts. Molecular Ecology *2*, 113–118.

Lundberg, D.S., Yourstone, S., Mieczkowski, P., Jones, C.D., and Dangl, J.L. (2013). Practical innovations for high-throughput amplicon sequencing. Nat Methods *10*, 999–1002.

Wallace, J., Laforest-Lapointe, I., and Kembel, S.W. (2018a). Variation in the leaf and root microbiome of sugar maple (Acer saccharum) at an elevational range limit. PeerJ *6*.

Wallace, J.G., Kremling, K.A., Kovar, L.L., and Buckler, E.S. (2018b). Quantitative Genetics of the Maize Leaf Microbiome. Phytobiomes Journal *2*, 208–224.

White, T.J., Bruns, T., Lee, S., and Taylor, J. (1990). Amplification and direct sequencing of ribosomal RNA genes for phylogenetics. In PCR Protocols, M.A. Innis, D.H. Gelfand, J.J. Sninsky, and T.J. White, eds. (San Diego: Academic Press), pp. 315–322.
